# Endothelial *Adgrl2* expression and alternative splicing controls the cerebrovasculature

**DOI:** 10.1101/2025.10.24.682442

**Authors:** Alexander King, Catherine Garcia, Crisylle Blanton, Anna Chen, Amna Ahmad, David Lukacsovich, Csaba Földy, Takako Makita, Garret R. Anderson

## Abstract

Central nervous system development requires parallel but interrelated processes of neural circuit assembly and vascularization. Intersecting between these two processes, the cell-adhesion G-protein coupled receptor *Adgrl2* is expressed in select neuron populations where it has been found to localize and control the assembly of specific synaptic sites. Further, *Adgrl2* is found to be expressed in select non-neuronal brain cells, where it is restricted to endothelial cells. Testing for *Adgrl2* function in these cells, here we find that endothelial cell specific *Adgrl2* deletion results in an impairment in cerebrovascular integrity. To understand how it might be possible for *Adgrl*2 to function independently in neuronal and endothelial contexts, we analyzed single-cell RNA sequencing datasets for differences in transcriptional identity between these cell classes. Doing so, we find that expression of *Adgrl2* mRNA is subject to robust cell type-specific alternative splicing, resulting in distinct isoforms being produced in neurons compared to endothelial cells. Investigating the functional implication of this cell type-specific alternative splicing, we find that forced expression of the neuronal isoform of *Adgrl2* in endothelial cells leads to alterations in cerebrovascular properties. Morphologically, endothelial cells with the non-native neuronal *Adgrl2* isoform induce the formation of ectopic glutamatergic synaptic contacts onto endothelial cells, indicating alterations in the cell-cell recognition process. Functionally, in direct contrast to endothelial *Adgrl2* deletion, this genetic expression switch instead enhances blood-brain barrier integrity. This overly restrictive cerebrovascular function results in dysregulation of blood to cerebrospinal fluid homeostasis, enlargement of brain ventricles, and a higher risk of hydrocephalus. As such, alternative splicing serves as a cell type specific mechanism that provides isoform specific *Adgrl2* for discerning functions controlling neural circuit assembly and cerebrovascular homeostasis.

## INTRODUCTION

Vertebrate species have similar levels of protein coding genes yet diverge extensively in their cellular and tissue patterns (Ponting, 2008). To accommodate the complexity of tissue development in varying species, increased levels of mRNA alternative splicing contribute to the diversity of genetically encoded protein isoforms that are found with evolution (Barbosa-Morais et al., 2012). At the tissue level, alternative splicing and protein isoform diversity is notably most complex in the nervous system with vertebrate evolution (Barbosa-Morais et al., 2012; Merkin et al., 2012; Yeo et al., 2004). As the complexity of brain tissue increases, correct development of the assembly of unique neural circuits necessitates a proportional increase in the molecular players involved. To support the need for protein isoform diversity, neurons use specialized alternative splicing machinery which is considered critical for brain development (Porter et al., 2018). Neuron-specific alternative splicing is mediated by RNA binding proteins that are selectively expressed by neuronal populations, and have been implicated as critical regulators for determining neural circuit connectivity and synapse specification (Traunmüller et al., 2016).

Nervous system development, with its complex synaptic connection patterns, requires precise matching and synaptic adhesion between presynaptic and postsynaptic neurons amidst a diverse array of neuronal and non-neuronal options. This process of neural circuit assembly relies heavily on cell-cell recognition and synaptic adhesion which are mediated by various synaptic cell-adhesion molecules. Among these, is the latrophilin family (gene symbols *Adgrl1-3*) of adhesion G-protein coupled receptors (aGPCRs). This family not only undergoes extensive alternative splicing, but has also been implicated as essential molecular players in guiding neuronal circuit development (Knierim et al., 2019; Wang et al., 2024). As prototypical members of the aGPCR family, latrophilin proteins possess two powerful functional features uniquely suited for this role: Extracellular adhesion and intracellular signaling. Latrophilin forms trans-cellular adhesion complexes with Teneurins (*Tenm1-4*) (Silva et al., 2011) and Fibronectin leucine-rich transmembrane proteins (*Flrt1-3*) (O’Sullivan et al., 2012), interactions that are influenced by alternative splicing (Boucard et al., 2014; Li et al., 2020, 2018; Wang et al., 2021). Likewise, intracellular alternative splicing has been found to affect latrophilin function, including influence on GPCR signaling pathway specificity (Ovando-Zambrano et al., 2019; Wang et al., 2024). In neurons, *Adgrl* genes have established cell-cell recognition functions that integrate attraction and repulsive cues that are necessary for neuronal migration, axon guidance, and synapse assembly (Anderson et al., 2017; Donohue et al., 2021; Pederick et al., 2023, 2021; Sando et al., 2019; Sando and Südhof, 2021; Toro et al., 2020). Beyond the nervous system, latrophilins also have non-neuronal roles that are necessary for embryogenesis and cardiovascular function (Anderson et al., 2017; Camillo et al., 2021; Doyle et al., 2006; Lee et al., 2022, 2021, 2019; Tanaka et al., 2024).

As brain tissue is composed of both neuronal and non-neuronal cells that express *Adgrl* genes, we set out to understand the uniqueness of latrophilin functions in these distinct cell types. In this study, we investigate latrophilin expression and alternative splicing differences that exist amongst cell types. Doing so, we find a common theme that exists amongst the latrophilin family. While there are alternative splicing differences of *Adgrl* transcripts that are found between defined neuronal cell types, the greatest degree of splicing contrast is that observed between neurons and non-neuronal cells. Analyzing the variability in expression and alternative splicing, we find one gene to be most polarizing in both of these features, *Adgrl2*. Previously implicated as a key molecular player in neurons for controlling the patterning of neural circuits during synaptic assembly during development (Anderson et al., 2017; Donohue et al., 2021; Pederick et al., 2021), *Adgrl2* is also expressed by non-neuronal endothelial cells in the brain vasculature. Furthermore, we find that neurons and endothelial cells express uniquely different isoforms of *Adgrl2*. By expressing these distinct isoforms of *Adgrl2* in neurons and endothelial cells, we reveal a mechanism by which latrophilins serve a dual function in the intertwined processes of neural circuit formation and cerebrovascular development.

## RESULTS

### *Adgrl2* is selectively expressed amongst non-neuronal populations by endothelial cells and required for blood-brain barrier integrity

To further our understanding of cell type-specific expression and alternative splicing of latrophilin genes, we first set out to investigate the cellular and genetic profiles for the *Adgrl* family across large scale single-cell RNA sequencing (scRNAseq) datasets from the Allen Institute (Yao et al., 2021). Consisting of a robust collection of cortical and hippocampal cells (>73,000 cells; grouped into 42 subclasses based on similarities in their gene expression profiles using unsupervised clustering algorithms; see Supplemental Table 1), we surveyed the expression of *Adgrl* genes across both neuronal and non-neuronal cells. Examining their expression patterning, *Adgrl2* showed a cell type-specificity that was characteristically different from *Adgrl*1 or *Adgrl3,* which were more similar to each other (Figure 1A-C).

**Figure 1.**
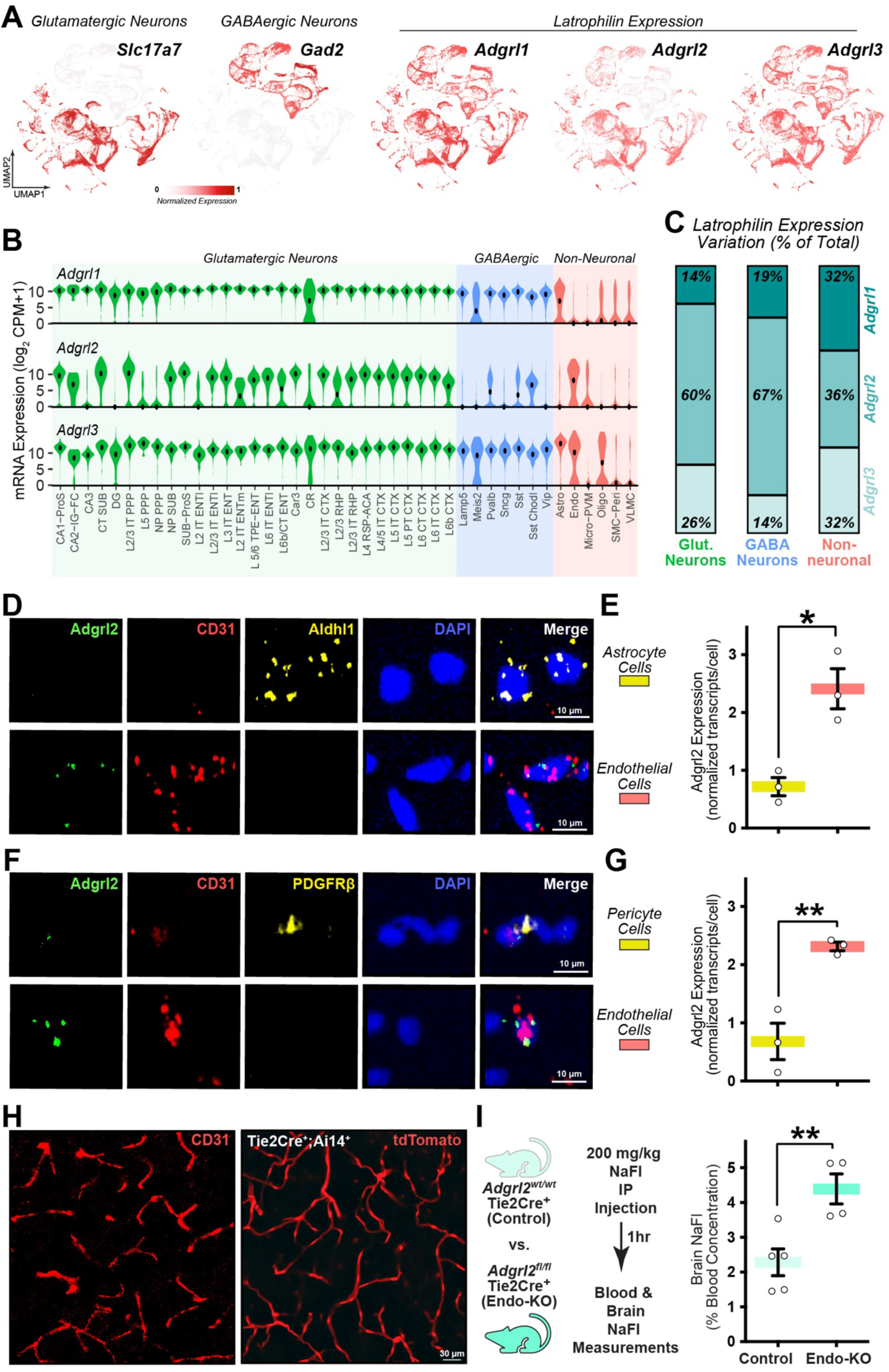
*Adgrl2* expression in endothelial cells is necessary for blood-brain barrier function. (**A**) UMAP representation of cortical and hippocampal cells for glutamatergic neuron marker *Slc17a7*, GABAergic neuron marker *Gad2*, and *Adgrl1-3* gene expression as published from the Allen Institute(Yao et al., 2021) (n = 73,348 cells). (**B**) Violin plots for *Adgrl1-3* expression within glutamatergic neuron (green), GABAergic neuron (blue), and non-neuronal (red) cell subclasses as indicated (see Supplemental Table 1 for expanded descriptions). Center dot represents median expression. (**C**) *Adgrl1-3* expression variation within glutamatergic, GABAergic, and non-neuronal cell classes. (**D**) Representative magnified cellular images of astrocytes visualized by Aldhl1 expression marker, and endothelial cells visualized by CD31 expression marker. (**E**) Quantitative analysis of *Adgrl2* expression in astrocytes and endothelial cells, with normalization and statistical methods described (n = 3 mice; P30-35). (**F**) Representative magnified cellular images of pericytes (tagged with Pdgfrβ) and endothelial cells (tagged with CD31). (**G**) Quantitative analysis of *Adgrl2* expression in pericytes and endothelial cells, with normalization and statistical methods described (n = 3 mice; P30-35). (**H**) Visualization of mouse brain vasculature using CD31 immunohistochemistry (left) and Tie2Cre-driven Ai14 reporter tdTomato expression (right). (**I**) Quantitative analysis of blood-brain barrier permeability between control (*Adgrl2*^wt/wt^;Tie2Cre^+^; n = 5mice; P30-35) and endothelial cell-specific *Adgrl2* knockout (Endo-KO) mice (*Adgrl2*^fl/fl^;Tie2Cre^+^; n = 4 mice; P30-35) using sodium fluorescein (NaFl) intraperitoneal (IP) injection and subsequent concentration measurements in blood and brain tissue. Data shown are means ± SEM. Statistical analysis was performed by a Student’s t-test (*p < 0.05, **p < 0.01).

To quantify this observed cell type specific expression, we used the metric tau (t) (Kryuchkova-Mostacci and Robinson-Rechavi, 2017; Yanai et al., 2005) to measure the variation in mRNA levels across cell subclasses. Doing so, we observed *Adgrl2* to possess the greatest amount of expression variation both in glutamatergic (∼60% of total *Adgrl* expression variation) and GABAergic (67% of total variation) neurons (Figure 1C). In non-neuronal cells, all *Adgrl* genes appeared to be restricted in their expression profile with comparable expression variations (Figure 1C). *Adgrl1* was selectively expressed by non-neuronal astrocytes, *Adgrl2* was largely restricted to endothelial populations, whereas *Adgrl3* showed variable expression across astrocytes, endothelial cells and oligodendrocytes (Figure 1B).

The extremes observed in cell type dependent genetic expression of *Adgrl2* isoforms suggest that *Adgrl2* has specialized roles that differentiate it from the other *Adgrl* genes. To investigate this further, we first sought to confirm its cell type-specific expression profile. Further, while *Adgrl* expression and function in neurons have been implicated in numerous neurodevelopmental processes including cell migration, axon guidance, and synaptic recognition and assembly (Anderson et al., 2017; Pederick et al., 2021; Sando et al., 2019; Toro et al., 2020), its function in non-neuronal cells in the brain has not been explored. Therefore, we set out to investigate the physiological importance of *Adgrl2* cell type specific expression in non-neuronal cells. To verify *Adgrl2* selective expression in endothelial cells amongst the non-neuronal population (Figure 1B), we assessed its expression through single molecule fluorescent in-situ hybridization (smFISH). Using smFISH probes against *Adgrl2* and endothelial specific transcript CD31 (Newman et al., 1990), we surveyed brains from mature (P30-35) mice to observe *Adgrl2* mRNA expression. Analyzing hippocampal containing brain sections where *Adgrl2* previously was found to be selectively expressed by CA1 pyramidal neurons (Donohue et al., 2021; Murphy et al., 2024; Pederick et al., 2021), we consistently found enriched *Adgrl2* expression in CA1 neurons and not detectable in the dentate gyrus granular neurons (Figure S1). Similarly, analyzing CD31+ cells in these same hippocampal slices, we find that *Adgrl2* is expressed in endothelial cells. In comparison to the nearby non-neuronal oligodendrocytes that are positioned in the alveus white matter tract, we find *Adgrl2* to be expressed at higher levels in CD31+ endothelial cells (Figure S1). To confirm the enrichment of *Adgrl2* expression in endothelial cells, we further examined its selectivity amongst other brain vasculature cell types. The cerebrovasculature is made up of not only endothelial cells but is supported by pericyte and astrocyte non-neuronal cells (Vanlandewijck et al., 2018). To survey *Adgrl2* expression amongst these vascular cell types, we performed smFISH analysis with multiple cell type marker probes. Using *Adgrl2* and CD31 probes, we included the probe Aldhl1 to also visualize astrocytes. Doing so, we find that *Adgrl2* was enriched in endothelial cells compared to astrocytes (Figure 1D-E). Similarly, using the pericyte marker Pdgfrβ in combination with *Adgrl2* and CD31 endothelial markers, we again find that *Adgrl2* was enriched in endothelial cells compared to pericytes (Figure 1F-G). Taken together, these data confirm that *Adgrl2* is expressed maximally in endothelial cells amongst the cerebrovascular cell types.

In addition to its role in nervous system development, *Adgrl2* has emerged as an important signaling molecule within the cardiovascular system. During embryonic heart development, *Adgrl2* has been shown to be a central signaling protein, as well as a marker for the differentiation of pluripotent stem cells into cardiac progenitor cells (Lee et al., 2022, 2021, 2019). In endothelial cells, *Adgrl2* has been implicated to control peripheral vascular permeability in zebrafish models by regulating endothelial tight junctions (Camillo et al., 2021). The impact of *Adgrl2* deletion on brain vasculature, however, has not been explored. This prompted the question: Is *Adgrl2* endothelial expression also required for cerebrovascular integrity? To address this, we looked to develop a genetic mouse model to selectively delete *Adgrl2* from endothelial cells. To do so, we genetically crossed *Adgrl2* conditional knockout mice (*Adgrl2^fl^*) (Anderson et al., 2017) with transgenic mice expressing Cre-recombinase under the control of the endothelial specific promoter Tie2 (Tie2-Cre) (Kisanuki et al., 2001). To confirm the selectivity of Cre-recombinase expression, we first crossed Tie2-Cre mice with the mouse reporter line with Cre-recombinase dependent expression of tdTomato fluorescent protein (Ai14) (Madisen et al., 2010). Doing so, we found Tie2-Cre^+^;Ai14^+^ mice results in brain tissue with tdTomato expression found in the vasculature, and closely resembles that of immunohistochemical staining against the endothelial protein CD31 (Figure 1H). As such, we proceeded to genetically cross Tie2-Cre mice with *Adgrl2^fl^* mice to produce control (Tie2Cre^+^;*Adgrl2*^wt/wt^) and *Adgrl2* endothelial knockout animals (Tie2Cre^+^;*Adgrl2*^fl/fl^) (Figure 1I). Using the small molecule fluorescent tracer sodium fluorescein (NaFl; 0.376 kDa), we then performed intraperitoneal NaFl injections and assayed levels in blood and in brain tissue to test for cerebrovascular blood-brain barrier (BBB) integrity (Figure 1I). Comparing BBB permeability between Tie2Cre^+^;*Adgrl2*^wt/wt^ versus Tie2Cre^+^;*Adgrl2*^fl/fl^ mice, we find that targeted *Adgrl2* endothelial deletion results in increased NaFl penetration into the brain (Figure 1I). Thus, *Adgrl2* expression in brain endothelial cells appears to be essential for cerebrovasculature function.

### *Adgrl2* cell type specific expression in brain endothelial cells is conserved in the human cerebrovasculature

To validate the importance of *Adgrl2* non-neuronal cell type specific expression, we next sought to confirm its expression in the human brain cerebrovasculature. To do so, we assessed its expression from previously published single nuclei transcriptomes of cerebrovascular cells from post-mortem tissues pooled across six brain regions (Sun et al., 2023). This results in a single cell gene expression atlas of 22,514 cerebrovascular cells including endothelial, pericyte, smooth muscle, perivascular fibroblasts and ependymal cells (Figure 2A). Surveying *Adgrl1-3* expression amongst these cell types (Figure 2B), *Adgrl2* is found consistently enriched amongst the endothelial cell population (Figure 2C). *Adgrl3* expression in comparison, is mixed and found most highly expressed in pericytes, smooth muscle cells, and to a lesser extent in endothelial cells. *Adgrl1* is largely undetectable throughout these vasculature cell types. Focusing on endothelial cells in greater detail, there are transcriptomic signatures that allow for the identification of endothelial subtypes including arterial (aEndo), capillary (cEndo), and venule (vEndo) endothelial cells (Garcia et al., 2022; Yang et al., 2022). Surveying *Adgrl1-3* expression amongst these endothelial subtypes, *Adgrl2* is found at highest abundance amongst the latrophilin family in all subtypes, followed by less abundant *Adgrl3*, and with *Adgrl1* largely undetected (Figure 2D). The expression levels also correlate with detection probability, with *Adgrl2* expressing cells consistently outweighing other latrophilin gene expression for all three endothelial classes including aEndo (∼37% *Adgrl2+* vs. ∼15% A*dgrl3+* vs. ∼8% *Adgrl1+*), cEndo (∼59% *Adgrl2+* vs. ∼26% *Adgrl3+* vs. ∼9% *Adgrl1+),* and vEndo cells (∼76% *Adgrl2+* vs. 60% *Adgrl3+* vs. ∼13% *Adgrl1+)* (Figure 2E). Taken together, *Adgrl2* expression enrichment in endothelial cells appears to be a conserved feature in mammalian evolution.

**Figure 2.**
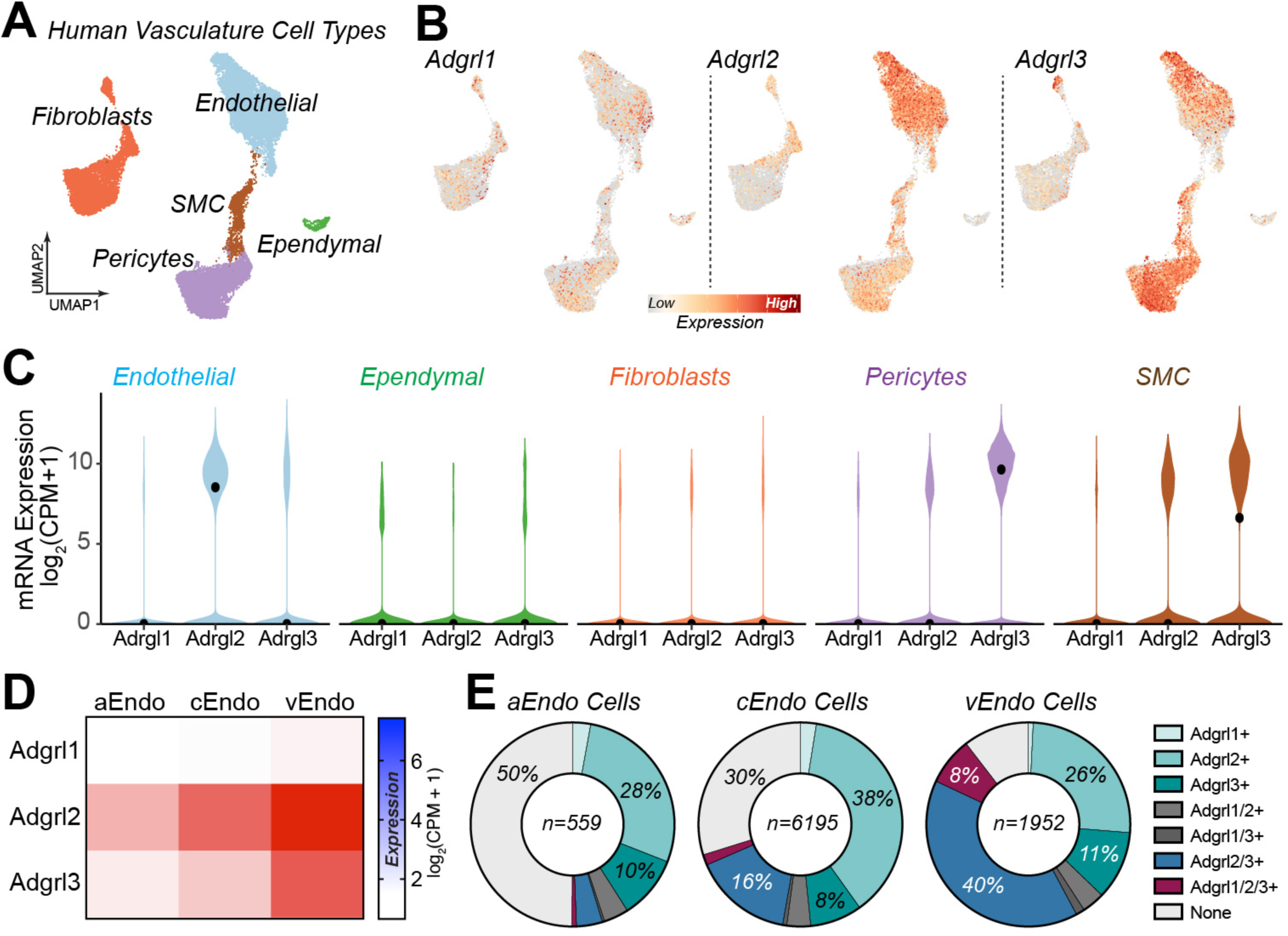
Enriched *Adgrl2* expression in endothelial cells is conserved in human cerebrovasculature cell types. (**A-E**) Analysis of human cerebrovasculature cell types from postmortem tissues as previously published (Sun et al., 2023). (**A**) UMAP representation of human cerebrovasculature cell types (endothelial, ependymal, fibroblasts, pericytes, and smooth muscle cells (SMC)) labelled according to their identifiable transcriptional signatures (n = 22,514 cells; 428 individuals). (**B**) *Adgrl1-3* gene expression within UMAP representations. (**C**) Violin plots of *Adgrl1-3* expression levels across the indicated vasculature associated cell types. Black dots indicate median expression level (n = 8706 endothelial, 340 ependymal, 6178 fibroblasts, 6004 pericytes, 1286 SMC). (**D**) Heat map *Adgrl1-3* expression plots for transcriptionally identified arterial (aEndo; n = 559 cells), capillary (cEndo; n = 6195 cells), and venous (vEndo; n=1952 cells) subtypes of endothelial cells. (**E**) Percentage of aEndo, cEndo, vEndo cells with detectable latrophilin expression (*Adgrl+*) categories as indicated.

### *Adgrl2* transcripts are alternative spliced in neurons versus endothelial cells

Given that *Adgrl* transcripts have previously been found to undergo extensive alternative splicing (Knierim et al., 2019; Wang et al., 2024), we next looked at isoform variant expression within neuronal and non-neuronal cell classes (Figure 3). Using the Allen Institute Mouse Brain Smart-Seq scRNAseq dataset (Yao et al., 2021), we performed bioinformatic characterization of the alternatively spliced exons within the protein coding regions of *Adgrl1-3*, including rates of exon inclusion/exclusion and shifts in exon junction sites (Figure 3A). Surveying known *Adgrl* exons (see Figure S2 and Supplemental Tables 2-4), we analyzed the exon inclusion proportion (EIP) across all *Adgrl1-3* scRNAseq reads that cover splice junctions (enabling to determine the proportion of reads that include or exclude a specific exon). Examining the EIP variation that exists among cell types (Figure 3B), a similar pattern emerges to that observed in expression variation (Figure 1C). *Adgrl2* (54%) exhibits the greatest amount of splicing variation amongst the latrophilin family, followed by *Adgrl3* (34%) and *Adgrl1* (12%). To examine these general trends within *Adgrl* splicing variation, we first looked at *Adgrl* EIPs across three broad cell classifications: glutamatergic (or excitatory) neurons, GABAergic (or inhibitory) neurons, and non-neuronal cells. With this analysis we find that *Adgrl1* exhibits limited alternative splicing that is confined to largely one major exon (7b) (Figure 3C-D). With *Adgrl2* and *Adgrl3* on the other hand, more complex alternative splicing patterns emerge with multiple coding exons being impacted. For *Adgrl2* we detect four major exons (9, 14, 23, 28) that exhibit large variations in EIP detection, and at 6 additional exons (11, 24, 27, 29-31) with lower levels of variation (Figure 3E-F). Similarly, *Adgrl3* exhibits splicing variations at four exons (exons 6, 9, 15, 24) and five additional exons that encode for the C-terminal region (exons 29-32) (Figure 3G-H).

**Figure 3.**
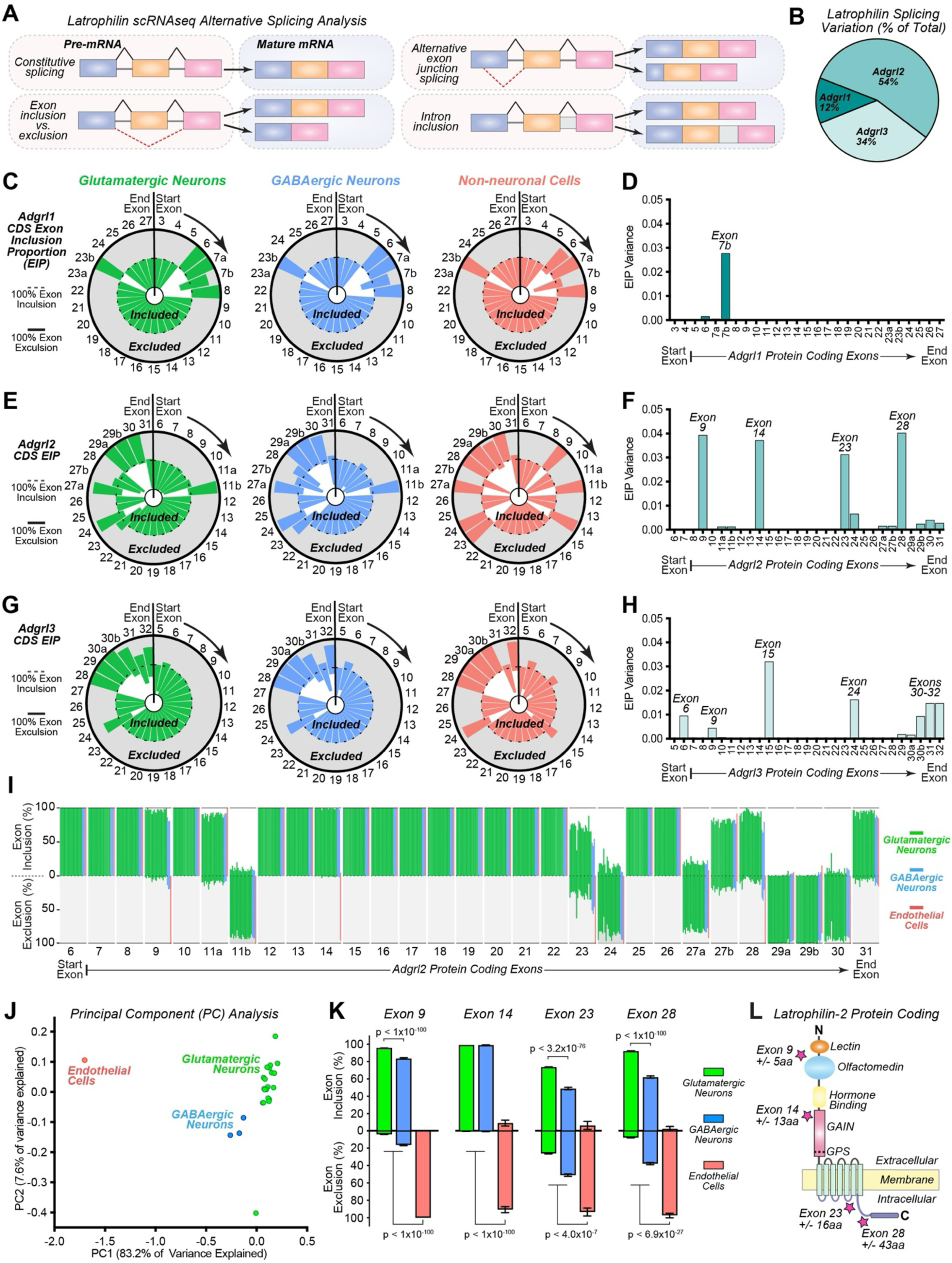
*Adgrl2* alternative splicing is cell class dependent at four major structural regions. **(A)** Illustration of alternative RNA splicing possibilities, examined for *Adgrl* splice variants. Alternative splicing events considered include exon skipping, alternative 5’ and 3’ splice sites, and mutually exclusive exons. (**B**) *Adgrl1-3* mRNA splicing variation, as measured by the variance (standard deviation squared) of exon inclusion proportion (EIP) across all cell subclasses transcriptomes previously published by the Allen Institute (Yao et al., 2021). (**C-D**) EIP analysis of protein coding exons (see Figure S2) for *Adgrl1*. (**C**) Shown are EIP circle charts with 5’ start exon indicated, and downstream exons progressing clockwise. Indicated exon is maximally included if graphic slice is confined to interior circle (dashed line) or maximally excluded if slice is restricted to shaded gray region of outer circle (solid line). (**D**) EIP variance between cell subclasses, as quantified by individually defined exons. (**E-F**) Similar as described for C-D, except for *Adgrl2*. (**G-H**) Similar as described for C-D, except for *Adgrl3*. (**I**) Summary graph of exon inclusion proportions (top) and exon exclusion proportions (bottom) across cell subclasses with detectable *Adgrl2* expression across the protein coding exons. (**J**) Principal component analysis (PCA) for different cell subtypes based on their alternative splicing patterns. Each point corresponds to a cell subtypes color coded by major classifications including glutamatergic (green), GABAergic (blue), and non-neuronal cells (red). Plot axes representing the first two principal components that capture the majority of variance in splicing profiles. (**K**) Summary graphs of exon inclusion proportions (top) and exon exclusion proportions (bottom) for the major alternatively spliced exons (9, 14, 23, and 28) averaged across all Glutamatergic and GABAergic neurons, in comparison to endothelial cells. Statistical analysis was performed by Dunn’s Test of multiple comparisons, with significance indicated. Only those cells containing transcripts that detected inclusion or exclusion of an exon were included for statistical analysis (n = 12008-24713 Glutamatergic neurons; 1229-3156 GABAergic neurons; 30-63 non-neuronal cells). (**L**) Visualization of *Adgrl2* protein structure. Stars indicate the domain regions where exons 9, 14, 23, and 28 alternative splice sites would impact protein coding.

With *Adgrl2* transcripts found to exhibit large variations in cellular alternative splicing patterns, to examine this in greater detail we next surveyed its splicing within defined cellular subclasses (Figure 3I). Breaking down *Adgrl2* exon inclusion proportions by cell types that have comprehensive exon detection, we performed principal component analysis (PCA) to visualize the EIP variation that exists between subclasses (Figure 3J). PCA of the *Adgrl2* EIPs variations across cell subclasses revealed clear separation between glutamatergic and GABAergic neuronal types, however the most robust differential splicing profiles are that found when comparing neuronal and endothelial cells (Figure 3J). When comparing Glutamatergic neurons, GABAergic neurons, and endothelial cells, we compared the splicing patterns at four main exons (9, 14, 23, 28) that exhibit the greatest variation (Figure 3K). Analysis of individual subclasses reveals that among the cells that prominently express *Adgrl2*, they generally follow the trends demonstrated in the PCA. Glutamatergic cells generally include their exons, endothelial cells tend to exclude them, and GABAergic cells tend to be intermediate; Exon 9 (EIP: ∼96% Glutamatergic vs. 84% GABAergic vs. 0% Endothelial), exon 14 (EIP: ∼99% Glutamatergic vs. 99% GABAergic vs. 10% Endothelial), exon 23 (EIP: ∼74% Glutamatergic vs. 49% GABAergic vs. 7% Endothelial), and Exon 28 (EIP: ∼92% Glutamatergic vs. 62% GABAergic vs. 3% Endothelial). Importantly, all four of these exons share one feature in common: the number of nucleotides in each of these exons is divisible by three. As such, inclusion of any of these exons would encode for a fixed amino acid insertion that impacts the immediate domain in the *Adgrl2* protein, without influencing the downstream sequences. Structurally, these exons (9/14/23/28) would impact protein encoding at four key structural regions (Figure 3L). Exon 9 is a mini-exon that encodes for 4 amino acids positioned between the lectin and olfactomedin extracellular domains, a region in the homologous *Adgrl1* exon that functionally impacts the affinity of latrophilin-1 interactions with teneurin extracellular binding partners (Boucard et al., 2014). Exon 14 encodes for 13 amino acids positioned within the GAIN domain, a structural region shared between all aGPCRs (Araç et al., 2012). Exon 23 encodes for 15 amino acids positioned within the third intracellular loop of the GPCR structure, a key region that determines G-protein signaling selectivity (Sadler et al., 2023). And lastly, exon 28 encodes for 43 amino acids positioned at the intracellular C-terminal region. Performing similar analysis for *Adgrl1* (Figure S3) we find Adgrl1 to be restricted in its alternatively splicing, confined to the homologue of *Adgrl2* exon 9. *Adgrl3* however, is similarly spliced at all four homologues of *Adgrl2* exons 9/14/23/28 (Figure S4). Of note, what is consistently found across all three latrophilin genes, the greatest difference in alternative splicing is that observed when comparing neuronal with non-neuronal cell types.

To validate neuronal versus endothelial alternative splicing of *Adgrl2* further, we looked to confirm these results against independent neuronal and endothelial RNAseq datasets (Figure S5). This includes neuronal RNAseq datasets from ribosome-associated mRNAs using large-scale tagged-ribosomal affinity purification (RiboTRAP) (Furlanis et al., 2019), neuronal scRNAseq datasets from acute slice embryonic neurons (Lukacsovich et al., 2019), and brain and peripheral endothelial specific scRNAseq datasets (Vanlandewijck et al., 2018). Doing so, we found consistency in neuronal *Adgrl2* splicing patterning outside of a few exons (14, 24, 27a/b) that appear developmentally regulated. Notably, exons 14 and 27b are included at a higher rate in adult glutamatergic and GABAergic cells in comparison to their embryonic counterparts (Figure S5). Conversely, exons 24 and 27a are included in higher proportions in embryonic cells in comparison to their adult counterparts. Yet, these differences are relatively subtle compared to endothelial *Adgrl2* alternative splicing profiles, which are consistently similar in both the brain and peripheral lung, regardless of the study dataset (Figure S5). Altogether, these results show that *Adgrl2* is alternatively spliced at four major exon positions, with the most robust cell type specific differences observed when comparing between glutamatergic neurons and endothelial cells. Glutamatergic neurons mostly include the major alternative spliced exons 9, 14, 23, and 28. In striking contrast however, endothelial cells mostly exclude these exons.

### Endothelial *Adgrl2* alternative splicing program prevents ectopic synaptogenesis

With endothelial *Adgrl2* expression now established to be essential for blood-brain barrier functionality, we next asked: What role does endothelial isoform specific expression of *Adgrl2* have for cerebrovasculature physiology? To answer this, we turned to a knock-in mouse model of *Adgrl2* (*Adgrl2^KI^*)*. Adgrl2^KI^*mice forces the expression of a single isoform of *Adgrl2*, under the control of the endogenous *Adgrl2* promoter (Anderson et al., 2017). This knock-in model expresses the neuronal isoform of *Adgrl2* (see NCBI reference sequence NP_001074767) in all cell-types that express *Adgrl2*. Thus, in endothelial cells that would ordinarily express the *Adgrl2* isoform lacking exons 9/14/23/28, *Adgrl2^KI^* endothelial cells express the isoform with their inclusion (Figure 4A).

**Figure 4.**
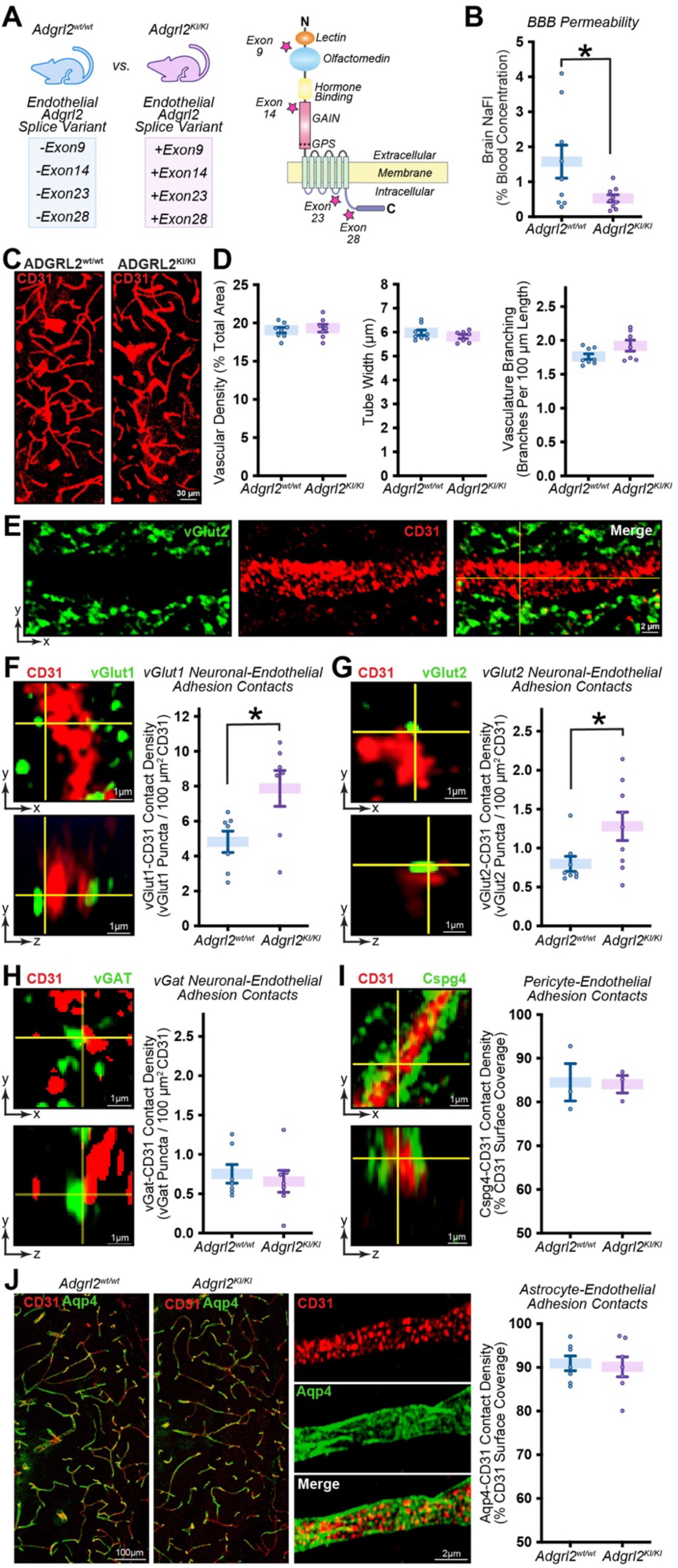
Expression of the glutamatergic neuronal isoform of *Adgrl2* in endothelial cells results in BBB permeability changes and ectopic synapse formation. (**A**) Hypothesis of *Adgrl2* isoform switching in endothelial cells. While wild-type mice express the *Adgrl2* isoform in endothelial cells that lack key alternatively spliced exons (-Exon9/14/23/28), the isoform specific *Adgrl2* knock-in line (*Adgrl2*^KI^) expresses the isoform with their inclusion (+Exon9/14/23/28). Shown is a schematic representation of the *Adgrl2* protein structure, with domains affected by alternative splicing. (**B**) Summary graph of BBB function assessed by NaFl permeability assay in wild-type (wt) and homozygous *Adgrl2*^KI/KI^ mice (n = 9 mice; P30-35). (**C**) Representative images of mouse brain vasculature in the hippocampal CA1 region visualized using CD31 immunohistochemistry. (**D**) Summary graphs of vascular architecture metrics including density (left), vessel widths (middle), and branch points (right) between wild-type (n = 8 mice; P30-35) and *Adgrl2*^KI/KI^ mice (n = 7 mice; P30-35). (**E**) Confocal image from hippocampal CA1 stratum lacunosum-moleculare (SLM) region, with immunohistochemistry labeled endothelial cells (CD31) co-labelled with the excitatory presynaptic marker vGlut2. (**F**) *Left,* Magnified images of example vGlut1 contact points with endothelial cells in the CA1-SLM region. *Right*, summary graphs of vGlut1 contact point densities on CD31 labeled endothelial surfaces in wild-type and homozygous *Adgrl2*^KI/KI^ mice (n = 7 mice; P30-35). (**G**) Similar as described for (F), but for vGlut2 contact points with endothelial cells (n = 8 mice; P30-35). (**H**) Similar as described for (F), but for vGat contact points with endothelial cells (n = 7 mice; P30-35). (**I**) *Left,* Magnified images of pericyte marker Cspg4 in contact with endothelial cells. *Right*, summary graphs of Cspg4 surface coverage of CD31 signal in wild-type and *Adgrl2*^KI/KI^ mice (n = 3 mice; P30-35). (**J**) *Left,* Overview and magnified images of astrocyte marker Aqp4 in contact with CD31. *Right,* summary graphs of Aqp4 surface coverage of CD31 signal in wild-type and *Adgrl2*^KI/KI^ mice (n = 7 mice; P30-35). Data shown are means ± SEM. Statistical analysis was performed by Student’s t-test (*p < 0.05).

With this transgenic mouse line, we next sought to investigate how this *Adgrl2* splicing manipulation impacts the cerebrovasculature. We first surveyed the impact on BBB function by comparing wild-type control mice (*Adgrl2^wt/wt^*) with homozygous *Adgrl2^KI/KI^* animals. Doing so, we find surprisingly, that *Adgrl2^KI/KI^* animals does not result in an impairment in BBB function, but rather exhibit an apparent enhancement in cerebrovascular integrity (Figure 4B). Thus, in contrast to the BBB impairment that is observed with *Adgrl2* deletion (Figure 1K), *Adgrl2*^KI/KI^ mice appear to lead to the opposite. To assess if this was a result of any cerebrovasculature morphological changes, we immunostained brain sections from *Adgrl2*^wt/wt^ and *Adgrl2^KI/KI^* animals for the endothelial cell marker CD31 (Figure 4C). However, after visualizing and quantifying the vasculature, we found no differences in gross morphological parameters including blood vessel density, width, and branching (Figure 4D). Having ruled out alterations in vascularization morphology to potentially explain the changes in BBB properties observed in *Adgrl2^KI/KI^* animals, we next sought to test the hypothesis that manipulation of endothelial *Adgrl2* isoform expression would impact the cell-adhesion recognition process of vasculature cells. Our results suggest that neurons express one version of *Adgrl2* (+Exons 9/14/23/28), which would be used to promote its synaptogenesis function in those cell types. Likewise, endothelial cells express an *Adgrl2* isoform that lacks these key exons (-Exons 9/14/23/28), which serves to differentiate it from neuronal recognition and permit instead for *Adgrl2* vasculature functions. Therefore, we postulate that forced expression of the neuronal version of *Adgrl2* in endothelial cells, would lead to a gain of function effect and promote synaptogenesis onto these cell types. To investigate this possibility, we looked to examine endothelial cells in a brain region with a well-established synaptogenesis role for *Adgrl2*. In the stratum lacunosum moleculare (SLM) region of the hippocampus, *Adgrl2* protein is found to be tightly coupled to excitatory synapses where it controls synaptogenesis from entorhinal cortical axonal projections (Anderson et al., 2017; Murphy et al., 2024; Sando et al., 2019). Focusing on this hippocampal SLM region, we performed immunohistochemical visualization of endothelial cells with the marker CD31, coupled with the glutamatergic presynaptic markers vGlut1 and vGlut2. In these experiments, vGlut and CD31 visualization is essentially mutually exclusive, with very little overlap between their signals (Figure 4E). The only sites of signal overlap are at sparse, but detectable contact points where vGlut is adjacent to CD31 marked endothelial cells (Figure 4F-G). While these are rare occurrences, we proceeded to quantify these neuronal-endothelial contact points. We found that compared to wild-type control mice, both vGlut1 and vGlut2 contact points with endothelial cells were enhanced (∼2-fold) in *Adgrl2^KI/KI^* animals (Figure 4F-G). Thus, endothelial cells possess a gain of function effect in *Adgrl2^KI/KI^* animals, resulting in increased excitatory synaptic contact points. To check the specificity of this observation, we next examined the impact on inhibitory synapses onto endothelial cells. Performing similar experiments, we visualized CD31 alongside the GABAergic presynaptic marker vGAT. Comparing wild-type controls with *Adgrl2^KI/KI^* animals in these experiments however, we observe no difference in vGAT to CD31 contact points (Figure 4H). We then proceeded to look at cell-adhesion between endothelial cells and important members of the blood-brain barrier, pericytes and astrocytes. Using the pericyte marker Cspg4 in combination with CD31 antibodies, we surveyed pericyte coverage of endothelial cells, finding no differences between genotypes (Figure 4I). To visualize astrocyte-endothelial interactions, we similarly used the astrocyte marker that localizes to the vasculature, Aqp4 (Figure 4J). Again, no differences were found between genotypes. Thus, forced expression of the neuronal *Adgrl2* isoform in endothelial cells, leads to a selective impact on endothelial cell-adhesion. Specifically, endothelial cells in *Adgrl2^KI/KI^* animals results in increased levels of neuronal synaptic contacts from glutamatergic neurons, with no impact on its adhesion with other cell types.

While there are gain of function effects found in *Adgrl2^KI/KI^* mice, deleterious effects on the cerebrovasculature are also observed. Notably, amongst homozygous *Adgrl2^KI/KI^* mice, we observed a modest yet distinct increase in the development of hydrocephalus neurological disorder (∼0.5% - 8 out of 1660 *Adgrl2^KI/KI^* 129S6xC57BL/6J hybrid animals; vs. 0 out of 680 *Adgrl2*^wt/wt^ 129S6xC57BL/6J hybrid animals). Hydrocephalus is a neurological condition characterized by an increase in cerebrospinal fluid (CSF) and brain ventricle volume (Zaksaite et al., 2023). This buildup can be the result of a number of causes including blocked CSF flow, increased CSF production, or reduced CSF absorption (Zaksaite et al., 2023). To investigate how manipulation of the endothelial *Adgrl2* isoform expression might lead to the development of hydrocephalus, we hypothesized that management of blood to CSF homeostasis is dysregulated in *Adgrl2^KI/KI^* mice. CSF is produced by specialized and highly vascularized choroid plexus tissue that is located in each of the ventricles of the brain(Lun et al., 2015). To confirm that *Adgrl2* is selectively expressed by endothelial cells within the choroid plexus, we surveyed *Adgrl2* expression with smFISH in combination with CD31 endothelial probe on the choroid plexus tissue positioned adjacent to the dorsal 3^rd^ ventricle (Figure 5A). Doing so, we find that CD31+ endothelial cells in the choroid plexus express *Adgrl2*, whereas epithelial cells that directly line the ventricle do not (Figure 5A-B). To validate this selective expression profile in the choroid plexus and its applicability to human physiology, we turned to an independent study with single cell transcriptomes isolated from postmortem human cortex and choroid plexus (Figure 5C-D)(Yang et al., 2021). Consistently, amongst this dataset, *Adgrl2* is found selectively expressed in endothelial cells amongst non-neuronal cells in both the cortex (Figure 5C) and choroid plexus (Figure 5D).

**Figure 5.**
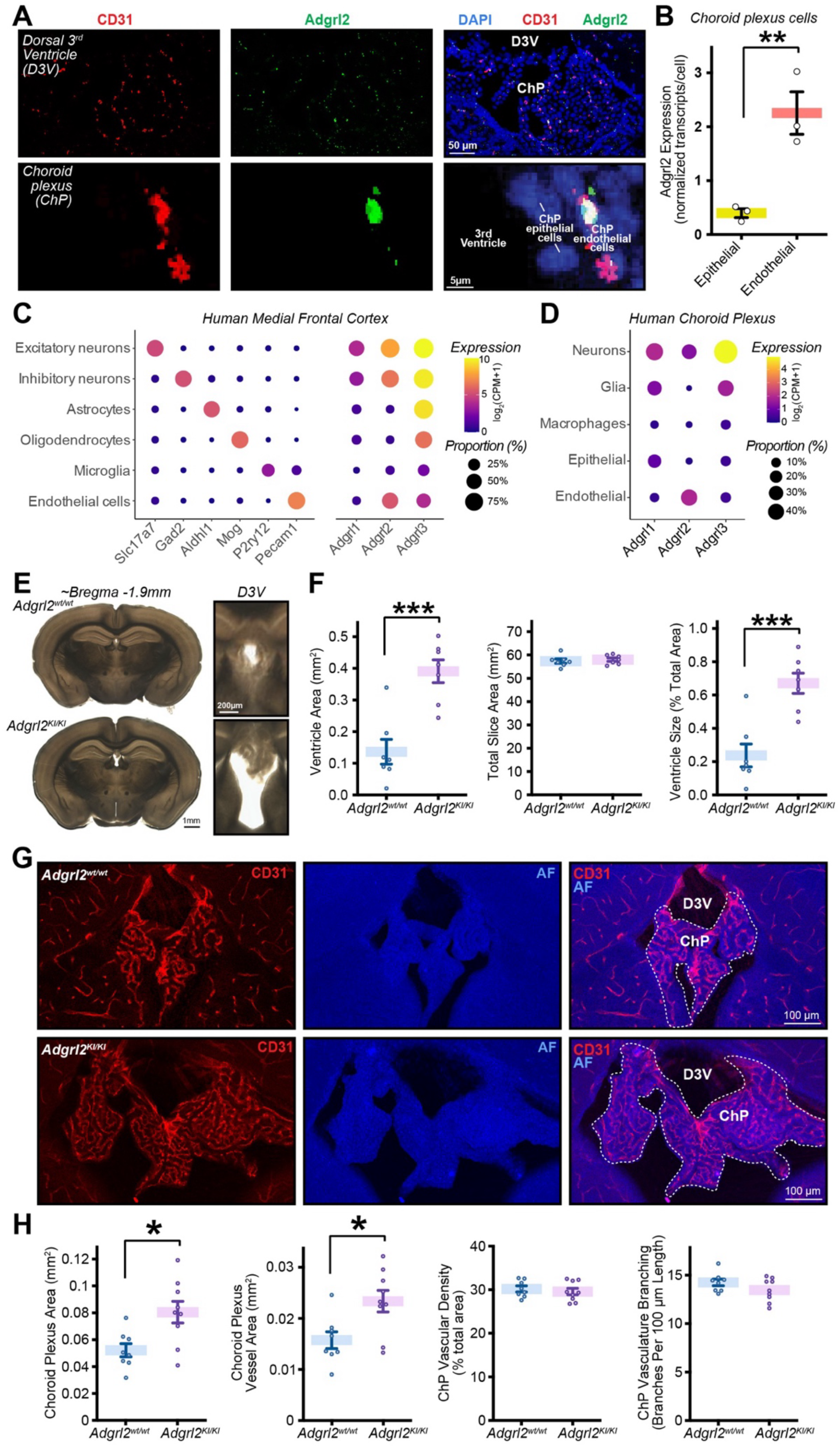
*Adgrl2* is selectively expressed by endothelial cells of the choroid plexus and required for normal ventricle development. (A) Representative confocal image of wild-type mice P30 coronal brain section (∼bregma −1.9 mm) of the dorsal third ventricle using single-molecule RNA in-situ hybridization for CD31 (red) and *Adgrl2* (green) mRNA. DAPI (blue) fluorescence shown for visualization of all nuclei. Shown are overview image (top) of the choroid plexus (ChP) tissue adjacent to the dorsal third ventricle (D3V), and zoomed in ChP cells adjacent to the D3V with epithelial cells and CD31+ endothelial cells indicated (bottom). **(B**) Summary graphs of *Adgrl2* expression for ChP epithelial and endothelial cells, normalized to average expression across the ChP (n = 3 wild-type mice; P30). (**C-D**) Analysis of *Adgrl1-3* expression levels in human postmortem tissue single cell transcriptomes isolated from the (**C**) Medial frontal cortex (38,217 cells from 16 individuals) or (**D**) Choroid plexus (27,092 cells from 14 individuals) (Yang et al., 2021). **(E**) Representative images of D3V containing coronal sections (P30-P37) from wild-type and *Adgrl2*^KI/KI^ mice. (**F**) Summary graphs of raw D3V area (*left*), total brain slice area (*middle*), and normalized D3V area (*right*) (n = 7 animals). (**G**) Representative ChP vascular visualization using CD31 (red) immunohistochemistry on D3V containing coronal sections (P30-P37) with tissue visualization by 405 nm autofluorescence (AF; blue) from wild-type and *Adgrl2*^KI/KI^ mice. (**H**) Summary graphs of ChP morphology including tissue size (*left*), vascular area (*middle left*), vascular density (*middle right*) and vascular branching patterns (*right*). Statistical analysis was performed by Student’s t-test (*p < 0.05, **p < 0.01, ***p < 0.001).

Since hydrocephalus is characterized by the development of enlarged ventricle space to accommodate increased CSF volume, we surveyed what impact this genetic manipulation has on ventricle size. Focusing on the dorsal 3^rd^ ventricle, in non-hydrocephalus *Adgrl2^KI/KI^* mice we observed an apparent ∼2.5-fold increase in ventricle size in comparison to wild-type mice (Figure 5E-F). We next surveyed the vasculature of the choroid plexus by IHC visualization with CD31 antibodies in both *Adgrl2*^wt/wt^ and *Adgrl2^KI/KI^* mice (Figure 5G). The choroid plexus possesses a distinct vascularization patterning that is not seen in other brain regions, making it readily identifiable. Surveying the morphology of the choroid plexus in the similar dorsal 3^rd^ ventricle, we also find a comparable increase in the size of the choroid plexus tissue (Figure 5H). This increase in choroid plexus tissue appears to be fully vascularized as a proportional increase in endothelial vessel staining is observed, with no apparent change in other morphological parameters including vascular density and branching patterns (Figure 5H). Therefore, both an increase in choroid plexus tissue and a correlated increase in ventricle size is observed in *Adgrl2^KI/KI^* mice, suggesting increased production of CSF as a likely contributing factor to the higher prevalence of hydrocephalus.

Altogether, forced expression of the neuronal isoform of *Adgrl2* in endothelial cells leads to both gain of function, and loss of function changes in the cerebrovasculature. Gain of function effects include an increase in glutamatergic presynaptic sites in contact with endothelial cells, and an increase in the restrictive properties of the blood-brain barrier. Loss of function effects include dysregulation of normal blood to cerebrospinal fluid homeostasis, resulting in brain ventricle enlargement and a higher likelihood in the development of hydrocephalus neurological disorder.

## DISCUSSION

The development of the mammalian brain requires the formation and proper function of two major parallel yet interdependent networks: neural circuitry and vascularization. While clearly distinct, the development of these networks relies on genetically programmed mechanisms that share common molecular signaling pathways and organizational principles. The concept known as the “neurovascular link,” which describes the resemblance between vascular and neural growth, was initially observed in the peripheral nervous system (Carmeliet and Tessier-Lavigne, 2005). However, it has since been discovered that this principle also extends to the central nervous system, where angiogenesis and vascularization depend on molecular cues that are shared with neurons (Wälchli et al., 2015). These shared molecular players and signaling pathways play pivotal roles in various processes including axon growth and guidance during neural circuit development, as well as for angiogenesis of embryonic and postnatal vascularization of the brain. Neural circuit development requires complex repulsion and adhesion between specific cell types to control numerous processes including cell migration, neural dendritic and axonal growth, synaptic target recognition, and synaptogenesis. Similarly, the cerebrovasculature requires tight endothelial-endothelial junction interactions, as well as endothelial interactions with pericyte and astrocyte cell types to support a functional blood-brain barrier. This study introduces a key molecular player that is shared between neural circuit development and the cerebrovasculature, *Adgrl2*.

During our expression analysis across defined brain cell types, we find a common mechanistic theme that is shared amongst the latrophilin genetic family. To differentiate neuronal versus non-neuronal *Adgrl* functions, these distinct cell populations alternatively splice *Adgrl* mRNA transcripts that results in the translation of distinct protein isoforms. The complexity of these alternative splicing patterns, however, is not shared across all *Adgrl* genes. *Adgrl1* is alternatively spliced at a singular extracellular site, whereas *Adgrl2* and *Adgrl3* contain multiple extracellular and intracellular regions subject to cell type specific alternative splicing. Among the *Adgrl* genes, *Adgrl2* shows the most pronounced differences in alternative splicing patterns across cell types. Where these differences are applicable however, is restricted to very specific cell populations. *Adgrl2* is expressed mainly in neuronal cell populations, yet is confined to select glutamatergic and GABAergic cell types. While these neuronal cells demonstrate detectable differences in their *Adgrl2* alternative splicing patterns, these differentiations are relatively subtle in comparison to endothelial cells that express *Adgrl2.* Comparing *Adgrl2* expressing neuronal and endothelial cell populations, we find transcripts to be alternatively spliced at four key exons. While neurons largely include these exons, endothelial cells excise them. This leads to cell type-specific transcripts of *Adgrl2* that differentially encode protein at four distinct domain regions. These include the extracellular lectin domain (exon 9), the extracellular GAIN domain (exon 14), the third intracellular GPCR loop (exon 23), and the intracellular C-terminal region (exon 28). As a result, endothelial and neuronal cells express different isoforms of *Adgrl2,* each with distinct functional properties. These variations can influence the specificity of *Adgrl2*-dependent extracellular adhesion interactions, GPCR signaling properties, and the targeting of intracellular trafficking to specific cellular locations.

Although the specific splicing factors responsible for cell-type-specific alternative splicing of *Adgrl2* remain unknown, it is well established that neurons and non-neuronal cells employ distinct mechanisms to control these processes (Mauger and Scheiffele, 2017). A key component of this machinery are RNA-binding proteins, which contain defined RNA-binding domains that recognize RNA sequences or structures to promote or inhibit exon inclusion (Tao et al., 2024). To distinguish neurons from other cell types, neuron-specific splicing regulators are necessary components of the alternative splicing machinery. Notable neuron-specific splicing factors include those that promote exon inclusion or exclusion such as the *PTBP* family (Keppetipola et al., 2012), *Nova* (Jensen et al., 2000), *RBFOX* (Jin et al., 2003; Weyn-Vanhentenryck et al., 2014), and *nSR100* (Irimia et al., 2014). To deepen our understanding of the mechanisms that enable cell type specific proteomes, future studies will need to identify which of these specific splicing factors are involved in controlling the differential latrophilin splicing that exists between neurons and non-neuronal cells.

To test the physiological relevance of latrophilin expression and alternative splicing in non-neuronal cells, we examined the loss of function and gain of function effects of the gene *Adgrl2* in non-neuronal endothelial cell populations. Focusing on the role of *Adgrl2* in endothelial cells and cerebrovasculature physiology, we characterized a mouse model with genetic deletion versus a mouse model with forced expression of the neuronal spliced isoform. While *Adgrl2* endothelial deletion results in an impairment in BBB integrity, *Adgrl2^KI/KI^* mice in contrast results in an even more restrictive BBB. Investigating this observation in *Adgrl2^KI/KI^* mice further, we find that the genetic manipulation does not disrupt endothelial cell-adhesion with the main cerebrovasculature cell types (i.e. pericyte and astrocyte adhesion). Interestingly however, we do detect alterations in cell-adhesion that originate from relatively minor cell-type that contacts endothelial cells – that from excitatory neurons. *Adgrl2^KI/KI^*mice exhibit enhanced direct excitatory presynaptic contacts onto endothelial cells, suggesting that there is a potential increase in direct neuron to endothelial communication. Although the physiological consequences of this observation need further investigation, the concept of direct communication between neurons and vasculature is well-established. This includes the process of neurovascular coupling, which enables local neural activation to increase regional blood flow, thereby meeting the increased demand for nutrients and facilitating the removal of metabolic waste (Kaplan et al., 2020). While the role of neurotransmitter action on brain endothelial cells directly is not as well understood, there is evidence that endothelial cells express both ionotropic and metabotropic receptors to classic neurotransmitters including glutamate and GABA (Lu et al., 2017; Tyagi et al., 2007). Further, genetic manipulation of glutamate receptors in endothelial cells has been implicated in varying cerebrovascular alterations, including impaired brain vasculature development with endothelial specific genetic deletion of the NR1 NMDA subunit (Kim et al., 2022).

In terms of blood-brain barrier function, acute activation of glutamate NMDA receptors with pharmacological delivery has been implicated as a triggering mechanism that results in the breakdown in BBB integrity (Epping et al., 2022; Kim et al., 2022; Vazana et al., 2016). This glutamate mediated breakdown of the BBB, however, is perhaps only physiologically relevant when excess glutamate is neuronally released during periods of regional neural network hyperexcitability, such as that observed in epileptic patients with active seizures (Marchi et al., 2012). More subtle activation of glutamate receptors in endothelial cells on the other hand, is poorly understood. Our results open the possibility that glutamate release by neurons in close proximity to endothelial cells, can have a direct impact on cerebrovasculature functionality. Normally endothelial cells use a version of *Adgrl2* that is reduced in its synaptogenic activity, which minimizes direct neuronal glutamate release sites onto the cerebrovasculature. By instead forcing endothelial cells to express the neuronal isoform of *Adgrl2* however, leads to an increase in glutamate presynaptic terminals in contact with endothelial cells. While it is not clear what role increased glutamatergic neuron to endothelial contact points has towards the cerebrovasculature impacts observed in *Adgrl2^KI/KI^* mice, the role of neuronal synapses in direct contact with the vasculature warrants further investigation.

Alternatively, the cerebrovascular alterations observed in *Adgrl2^KI/KI^* mice may be independent of its extracellular adhesion impacts. Instead, influences on intracellular signaling within specific subcellular compartments may be a direct contributing factor. To better understand the contributions of extracellular and intracellular functions of *Adgrl2* in endothelial cells towards cerebrovascular physiology, will require careful dissection of its molecular organization. Towards this goal, evaluating the splice site specific contributions of *Adgrl2* in endothelial cells will be needed, including their impacts on adhesion complexes, subcellular localization, intracellular signaling properties, and ultimately, cerebrovascular physiology.

While the vascular morphology and function appear largely intact in *Adgrl2^KI/KI^* mice with the subtle impacts noted above, the most robust disruption we do observe is that on CSF regulation. Specifically, *Adgrl2^KI/KI^* mice exhibit increased choroid plexus tissue, enlarged brain ventricles, and a higher incidence of hydrocephalus neurological disorder. While the underlying mechanism for aberrant growth of choroid plexus tissue is currently unknown, selective impacts on choroid plexus physiology have been observed in clinical settings. The abnormal expansion of choroid plexus tissue in these animals resembles pathological features seen in patients with choroid plexus pathologies including benign choroid plexus papillomas, malignant choroid plexus carcinomas, or choroid plexus hyperplasia (Cox et al., 2016; Price et al., 2023). These pathologies are relatively rare, but what they share in common is similar to that observed in *Adgrl2^KI/KI^* mice including the development of enlarged choroid plexus, increased CSF production, and associated with the development of hydrocephalus. Effective treatments for these pathologies remain elusive, largely due to poor understanding of choroid plexus biology. Thus, further investigation of *Adgrl2* function in the choroid plexus is warranted, to further our knowledge of choroid plexus tissue growth and how CSF homeostasis is regulated.

## METHODS

### Animals

*Adgrl2^fl^* and *Adgrl2^KI^* mice used in this study were described previously (Anderson et al., 2017). The original mouse line for the generation of *Adgrl2^fl^* and *Adgrl2^KI^*mice is available through the Jackson Laboratory Mouse Repository for distribution (B6;129S6-Adgrl2tm/sud/J, JAX Stock number: 023401). Tie2-Cre mice (Kisanuki et al., 2001) (B6.Cg-Tg(Tek-cre)1Ywa/J; Jackson Laboratory Stock#008863) and Ai14 tdTomato Cre-reporter mice (Madisen et al., 2010) (B6.Cg-Gt(ROSA)26Sortm14(CAG-tdTomato)Hze/J; Jackson Laboratory Stock #007914) were also obtained from the Jackson Laboratory. Mice were weaned at 21 days of age and housed in groups of 2 to 5 on a 12-hour light/dark cycle with food and water ad libitum in the University of California, Riverside Animal Housing Facility. All procedures conformed to National Institutes of Health Guidelines for the Care and Use of Laboratory Mice and were approved by the University of California, Riverside Administrative Panel on Laboratory Animal Care, and Administrative Panel of Biosafety. Male and female mice were used for all experiments in approximately equal proportions, in gender-matched littermate pairs. No obvious differences were noted due to gender.

### Single molecule RNA fluorescent in situ hybridization (smFISH)

P30 wild-type mice were anesthetized with with isoflurane in order to be perfused transcardially with 20 mL ice-cold 1% diethyl pyrocarbonate (DEPC) treated and autoclaved PBS, followed by 10 mL 1% DEPC treated and autoclaved PBS with 4% PFA. The acquired brains were first placed in a sterile solution of 4% PFA in PBS for 24 hours, before being placed in a 10% sucrose in PBS overnight at 4C. Brains were immersed in sterile 20% sucrose in PBS to be cryoprotected. Once it was no longer floating, it was placed in sterile 30% sucrose in PBS for 2-3 days. Prior to cryosectioning, brains were stored at −80 °C. Cryosectioned brains (with 15 µm thickness) were placed onto HistoBond+M adhesive microscope slides (VWR 16004-406) with precision glass slide covers (Thorlabs CG15KH1). For single molecule in-situ hybridization, probes used include: Latrophilin-2 (Mm-Lphn2, 319341; ACDBio), CD31 (Pecam1, 316721-C3; ACDBio), Pdgfrβ (Pdgfrβ, 411381-C2), and Aldh1l1 (Aldh1l1, 405891-C2). Akoya fluorophores used for hybridization include Opal 540 (C1 channel [Lphn2], FP1494001KT; Akoya Biosciences), Opal 620 (C2 channel [CD31], FP1495001KT; Akoya Biosciences) and Opal 690 (C3 channel [Pdgfrβ, Aldh1l1], P1497001KT; Akoya Biosciences). A Zeiss 880 confocal microscope with Airyscan using the Plan Apochromat 10x/0.45 M27 air objective scanning set in airy fast “Flex’’ mode (0.7x Nyquist) at 2.17 ms/pixel was used for imaging. A 405 nm laser and BP 420-480 + BP 495-550 emission filter with BP 420-460 + LP 500 secondary beam splitter was used to image the DAPI. A 488 nm laser with BP 420-480 + BP 495-550 emission filter and LP 525 secondary beam splitter was used to image Lphn2 probes after being hybridized to Opal 540. A 561 nm laser with BP 420-480 + BP 495-620 emission filter and LP 570 secondary beam splitter was used to image CD31 probes after being hybridized to Opal 620. A 633 nm laser with BP 570-620 + LP 645 emission filter and LP 660 secondary beam splitter was used to image Aldh1l1 and Pdgfrβ probes after being hybridized to Opal 690. A 405 nm laser with BP 420-480 + BP 495-550 emission filter and BP 420-460 + LP 500 secondary beam splitter was used to image DAPI stained cell nuclei. All captured images were put through analysis using ImageJ (https://imagej.nih.gov/ij/). After identifying neuron subtypes as DAPI+ nuclei assembled within a single cell type-specific marker, puncta were counted using QuPath 0.2.7 (https://qupath.github.io/) (Bankhead et al., 2017). Cell subtypes were identified as DAPI+ nuclei unambiguously collocated with a single cell type-specific marker, and *Adgrl2* puncta within each cell group were quantified if found within 5μm of the nucleus using the sub cellular spot detection tool. Puncta counts per cell were normalized to the mean puncta count per cell in the set of all cells analyzed in an image.

### scRNA-seq Data Acquisition and Processing

scRNAseq splicing data were gathered from GSE:185862 (Yao et al., 2021) and the cell metadata was collected from the Allen Brain Institute smartSeqV4 RNAseq database (https://portal.brain-map.org/atlases-and-data/rnaseq/mouse-whole-cortex-and-hippocampus-smart-seq). Cell sequence data was downloaded programmatically using parallel-fastq-dump (https://github.com/rvalieris/parallel-fastq-dump). For ascertaining gene counts, data were gathered directly from the Allen Brain Institute smartSeqV4 RNAseq database. For each gene in each cell, counts for that particular gene for that particular cell were divided by the total number of reads in that cell and then multiplied by 1 million to get the counts of reads of that gene per million reads, or counts per million (CPM). Raw reads were pre-processed using Trimmomatic software (Bolger et al., 2014). Processed sequencing reads were then aligned to the Genome Reference Consortium reference transcriptome version GRCm39 with NCBI Mus musculus Annotation Release 109, using the STAR aligner (Dobin et al., 2013) with the following parameters: trimLeft = 10, minTailQuality = 15, minAverageQuality = 20 and minReadLength = 30. For each cell, we calculated the total number of unique genes detected with at least 1 mapped read, and the number of mapped reads. We then calculated the median and median absolute deviation of these 2 values across all cells. Cells that were more than 3 median absolute deviations below the median in either category were rejected. Splice Junction counts and reads were analyzed using Python 3 (https://www.python.org/downloads/). Cells were categorized into classes and subclasses using the Allen Brain Institute smartSeqV4 hierarchical classification categorizations from the dataset metadata. Additionally, scRNAseq reads were analyzed from datasets containing immature embryonic neurons (Lukacsovich et al., 2019), cell-class specific riboTRAP (Furlanis et al., 2019), and endothelial cells from brain and lung (Vanlandewijck et al., 2018). For these datasets, alignment parameters were kept the same as described above, with the following modifications: The maximum splice junction database overhangs allowable change with each dataset. For each read that spans a splice junction, the number of bases belonging to a particular exon has a minimum of 1 and a maximum of the read length minus one. This overhang must be manually adjusted for each individual dataset. For Lukacsovich et al., the sjdboverhang = 150. For Furlanis et al., the sjdboverhang = 100. For Vanlandewijck et al., the sjdboverhang = 42. Additionally, the alignment was set to single-end reads.

### Immunohistochemistry (IHC)

Mice were anesthetized and perfused transcardially with 30 mL phosphate buffered saline (PBS; 137mM NaCl, 2.7mM KCl, 10mM Na2HPO4, 1.8mM KH2PO4, pH 7.1) and 10 mL freshly prepared 4% paraformaldehyde (PFA) in PBS. Brains were postfixed in 4% PFA in PBS for 2 hours at 4C. Brains were briefly rinsed in PBS, mounted in agarose and 100 mm horizontal serial sections were collected using a Vibratome VT100S (Leica Biosystems). For immunofluorescence staining, sections were then washed in PBS for 5 min under gentle agitation followed by incubation for 1 hour in a blocking solution containing 10% goat serum (ab7481; Abcam) and 0.5% Triton X-100 in PBS. Subsequently, sections were transferred into PBS containing 1% goat serum, 0.01% Triton X-100, primary antibody, and incubated overnight at 4C on a nutating mixer. Sections were washed twice with PBS for 5 min each, then twice with PBS for 15 minutes each, then incubated in a solution of 1% goat serum, 0.01% Triton X-100, and secondary antibodies for 4 hours at 4C. Sections were washed again twice with PBS for 5 min each, then twice with PBS for 15 minutes each, and mounted on microscope slides (MS10UW; Thorlabs) using Vectashield Plus Antifade mounting media (Vector Laboratories; H-1000-10) and Precision glass slide covers (Thorlabs CG15KH1). The following primary antibodies were used: CD31 (1:1000; rat polyclonal; 553370; BD Pharmingen), Cspg4 (1:200; rabbit polyclonal; 55027-1-AP; Proteintech), Aqp4 (1:200; rabbit polyclonal; #59678T; Cell Signaling Technology), VGLUT1 (1:500; guinea pig polyclonal; 135-318; Synaptic Systems), vGlut2 (1:1000; guinea pig polyclonal; 135-404; Synaptic Systems), vGat (1:1000; rabbit polyclonal; 131-004; Synaptic Systems), GFP (1:500; Rabbit polyclonal; Cat# A-11122; Invitrogen). The following secondary polyclonal antibodies with fluorophores were used: goat anti-rabbit Alexa-Plus 488 (1:1000 A32723TR; Thermo Fisher Scientific), goat anti-rat Alexa 594 (1:1000; A11007; Thermo Fisher Scientific), goat anti-mouse Alexa-Plus 633 (1:1000; A-21052 Thermo Fisher Scientific), and goat anti-guinea pig Alexa-Plus 633 (1:1000; A21105; Thermo Fisher Scientific). A Zeiss 880 confocal microscope with airyscan using the Plan Apochromat 40x/1.2 water objective scanning set in airy fast “Opt’’ mode (1.0x Nyquist) at 2.57 µs/pixel was used for imaging. A 405 nm laser and BP 420-480 + BP 495-550 emission filter with BP 420-460 + LP 500 secondary beam splitter was used to image the DAPI. A 488 nm laser with a BP 495-550 + LP 570 emission filter was used to image Aqp4, Cspg4, GFP probes. A 561 nm laser with BP 420-480 + BP 495-620 emission filter was used to image CD31. A 633 nm laser with a BP 570-620 + LP 645 emission filter was used to image vGlut1, vGlut2, and vGat. Z-stack images were captured in increments of 0.20 µm between slices.

### Blood-brain barrier integrity assay

To assess the integrity of the blood-brain barrier, the injectable fluorescent small molecule sodium fluorescein (Thermo Fisher Scientific A11659.2) was used as described previously (Ahishali and Kaya, 2020). Mice were injected intraperitoneally with a phosphate-buffered saline (PBS) solution containing 25 mg/mL of sodium fluorescein, with the administered amount calculated to be 200 mg per kilogram of body weight. After an hour, mice were anesthetized and perfused transcardially with 30 mL PBS and 10 mL freshly prepared 4% PFA. Immediately before perfusion, mouse blood was extracted via syringe from the left ventricle and then weighed. Following perfusion, both livers and brains were extracted from the mouse and then weighed. Brain or liver tissue was placed into 500 µL PBS for fluorescence analysis. Tissues were then homogenized. 500 µL of 60% trichloroacetic acid in PBS were then added to each sample, vortexed for 2 minutes, and kept at 4° C for 30 minutes. Samples were vortexed again for 2 minutes and centrifuged for 10 minutes at 18,000 x g. Then, 200 µL 37.5% NaOH was added to each sample. Supernatants were then loaded onto a 384-well black-bottomed well plate. Sample fluorescence was recorded on a Spectramax iD5 spectrophotometer plate reader (Molecular Devices) equipped with an excitation wavelength of 490 nm with an excitation wavelength range of +/- 7.5 nm and an emission wavelength of 530 nm with an emission wavelength range of +/- 13.5 nm. Sample values were then compared to a linear standard curve (50 ng/mL to 125 µg/mg) of sodium fluorescein to obtain tissue NaFl concentration levels.

## QUANTIFICATION AND STATISTICAL ANALYSIS

### scRNA-seq analysis

scRNAseq gene and exon counts were analyzed from the Allen Brain Institute smartSeqV4 RNAseq database (GSE:185862). To standardize the gene counts, they were adjusted to reflect counts per million (CPM) for each cell. Following this, each CPM value was transformed by adding one and then taking the base 2 logarithm. For measuring gene expression specificity, the metric Tau (t) was used as described previously(Kryuchkova-Mostacci and Robinson-Rechavi, 2017; Yanai et al., 2005), calculated as follows: t = Σ _i=1_(1-y_i_), (n-1). Where [y_i_ =x_i_, max(x_i_)], x_i_ defined as the mean log_2_ CPM of the gene in cell subclass i, and n is the number of cell subclasses. To analyze alternative splicing data from scRNAseq reads, Exon inclusion proportion (EIP) was calculated for each cell: EIP = {*I_exon_*}, {I_exon_ + 2’E_exon_}, where *I*_exon_ is the number of splice junction reads in the cell that include the exon and *E*_*exon*_ is the number of splice junction reads in the cell that exclude the exon. For exons with alternative junction splice sites within an exon (indicated by letter following exon number; e.g. 11a, 11b), the EIP is calculated such that the combined EIP of the alternative forms do not exceed one, treating them as mutually exclusive. If an exon has two splice junction variants, *a* and *b*, we use the following formula: EIP*_exon variant a_* = {I*_junction a_*}, {I*_junction a_* + I*_junction b_* + E*_exon_*}. Where *I_exon_* is the number of splice junction reads in the cell that include the junction unique to exon variant *a*, and *I_exon_* is the number of splice junction reads in the cell that include the junction unique to exon variant *b*. For a cell type to be analyzed for alternative splicing, a minimum of 100 cells and all exons required a minimal detection (inclusion or exclusion) level of >1 log_2_(CPM+1).

### Immunohistochemistry (IHC) 3-dimensional analysis

Confocal Z-stack images of IHC experiments were analyzed with Imaris 9.6 3-dimension image analysis software. Images were preprocessed with a gaussian filter and z-stack normalization to correct for any intensity attenuation along the z-axis. CD31 marked endothelial cells were reconstructed using the “surfaces” creation tool. Surface smoothing was applied with a radius of 1.2 µm, and vessels were thresholded with background subtraction. For astrocyte (Aqp4) and pericyte (Cspg4) reconstructions, surfaces were created with a smoothing radius of 1.0 µm. Endothelial surface coverage analysis by pericytes or astrocytes was performed using the surface-surface colocalization plug-in, and quantified as: % Endothelial Coverage = 100*(colocalization surface area)÷(total endothelial surface area). For vGlut1, vGlut2, and vGat interaction with endothelial cells, presynaptic terminals were reconstructed using the ‘‘Spots’’ creation tool to detect puncta with expected X/Y/Z-axis diameters set constant for all experiments (0.6 µm X-axis; 0.6 µm Y-axis; 1.2 µm Z-axis). Puncta detection from background involved an initial Gaussian smoothing of the images to enhance spot features, followed by a quality-based thresholding for spot detection. Background subtraction was performed using an additional Gaussian smoothing step to estimate and remove the background signal. Puncta were marked as localized to blood vessels if the puncta center point was found to be within 1 µm of the blood vessel surface. Endothelial associated puncta were subsequently normalized to the total surface area of CD31 signal.

### Vascular morphology analysis

Quantitative analysis of vasculature was performed with a custom Python script (https://doi.org/10.5281/zenodo.15802164) that utilizes scikit-image(Walt et al., 2014), aicsimageio (https://github.com/AllenCellModeling/aicsimageio), and Shapely (https://github.com/shapely/shapely) Python packages. For each image, a region of interest (ROI) was demarcated using QuPath 0.2.7 (https://qupath.github.io/) (Bankhead et al., 2017). The CD31 (vessel) channel was pre-processed with a median filter and Contrast Limited Adaptive Histogram Equalization (CLAHE)(Pizer et al., 1987) to reduce noise and enhance local contrast. Vessel-like structures were then enhanced using a multi-scale Frangi filter(Frangi et al., 2006), and the resulting detected vessels were segmented using a local adaptive Gaussian threshold. The binary vessel mask was refined by removing objects smaller than a minimum width threshold (2.0 µm). The network was then skeletonized to calculate total vessel length and identify branch points, which were consolidated to ensure a minimum spatial separation (10 µm). Key metrics, including Vessel Area Fraction (VAF), mean vessel diameter, and branch point density (per 100 µm of vessel), were calculated within the ROI.

### Dimension Reduction

Dimension reduction analysis was performed using principal component analysis (PCA) with prcomp function in R (https://www.r-project.org/). Variables were normalized using the prcomp function centering, and normalization was performed without scaling. For each gene, cell subclasses were included in the analysis if they met quality control thresholds (25-75% trimmed mean log2 (CPM+1) > 1 expression levels; cell count >100).

## Statistical Analysis

All data analysis was performed using Prism 10 software (GraphPad) or rstatix package in R (Kassambara, 2023). Data are shown as mean ± SEM. Significance testing was performed using Wilcoxon rank-sum, Kruskal-Wallis test, Dunn’s test of multiple comparisons, or two-tailed Student’s t-test, as indicated. Statistically significant differences are indicated by asterisks (*p<0.05; **p<0.01; ***p<0.001).

## Acknowledgements.

This work was supported by a grant from the Whitehall Foundation (2018-08-01), Regents Faculty Development Grant from UCR Academic Senate, and Initial Complement Funds from the University of California, Riverside to G.R.A.

## Author Contributions

A.K. designed, performed, and analyzed most experiments with methodology conceptualization provided by D.L., C.F., T.M., and G.R.A. Experimental analysis performed by C.G., C.B., A.C., and A.A. Paper written by A.K. and G.R.A. with input from all the authors. Project conceptualization, supervision, administration and funding acquisition by G.R.A.

## Declarations of interests

The authors declare no competing interests.

## Data availability

Raw sequencing data used in this study are accessible from NCBI Gene Expression Omnibus database through accession numbers GSE185862, GSE133291, GSE121653, GSE98816, GSE99058 and GSE99235. Raw and analyzed data will be provided upon request. Any additional information required to reanalyze the data reported in this paper is available from the lead contact upon request.

## Code availability

Publicly available code used in this study are accessible as described. Unique code generated in this study for vascular analysis is available at Zenodo (https://doi.org/10.5281/zenodo.15802164).

---

**Supplemental Figure 1.**
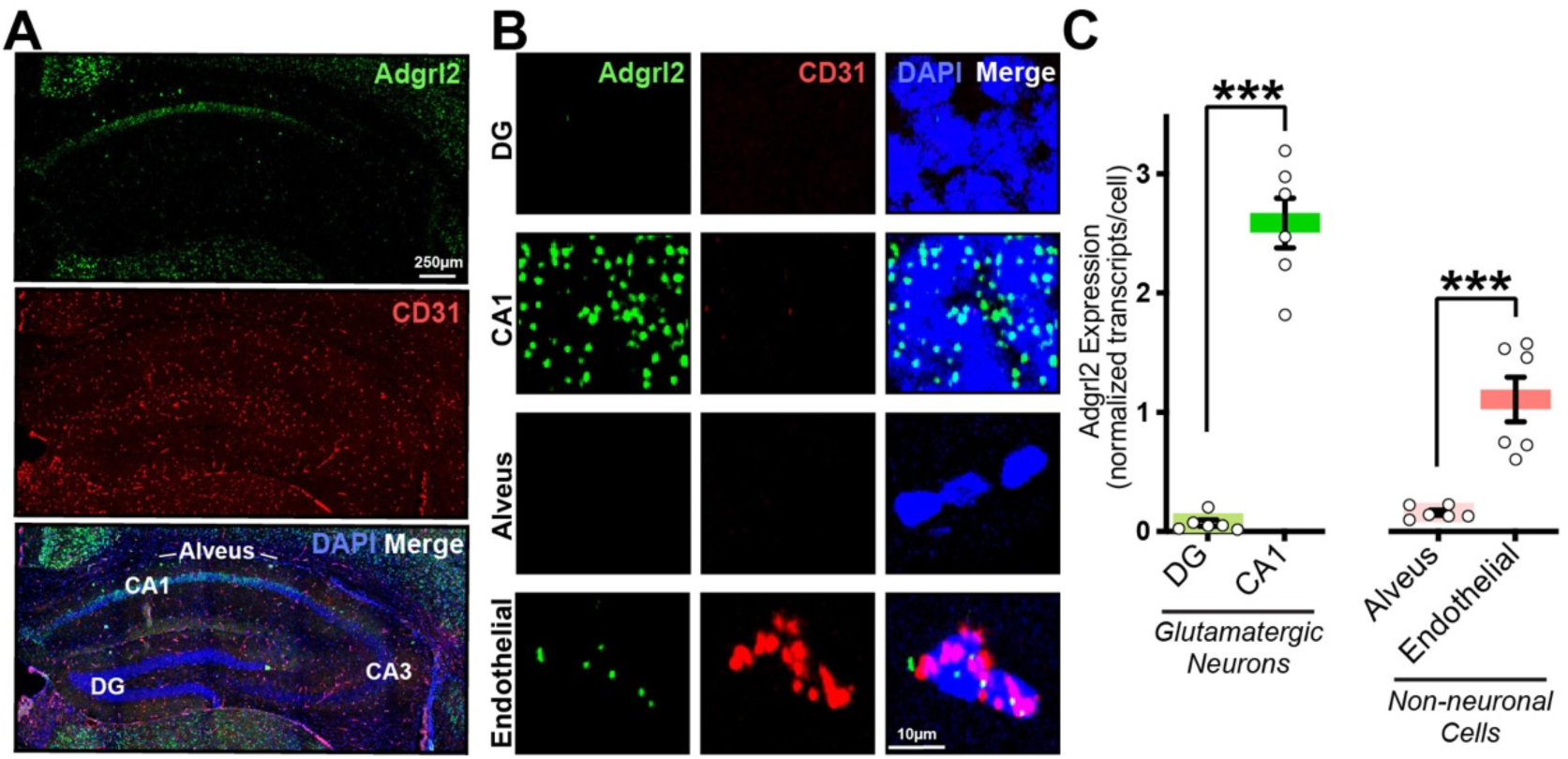
**(A**) Representative confocal image of P30 coronal brain section using single-molecule RNA in-situ hybridization for *Adgrl2* (green) and CD31 (red) mRNA. DAPI (blue) fluorescence shown for visualization of all nuclei. **(B**) Representative magnified cellular images of the CA1 pyramidal cell layer, the dentate gyrus granule layer, the alveus, and endothelial cells. (**C**) Summary graphs of *Adgrl2* expression in the CA1 pyramidal cell layer, dentate gyrus granule layer, alveus, and endothelial cells, normalized to the average expression across the hippocampus (n = 6 mice; P30-35). Data shown are means ± SEM. Statistical analysis was performed by a Student’s t-test (***p < 0.001).

**Supplemental Figure 2.**
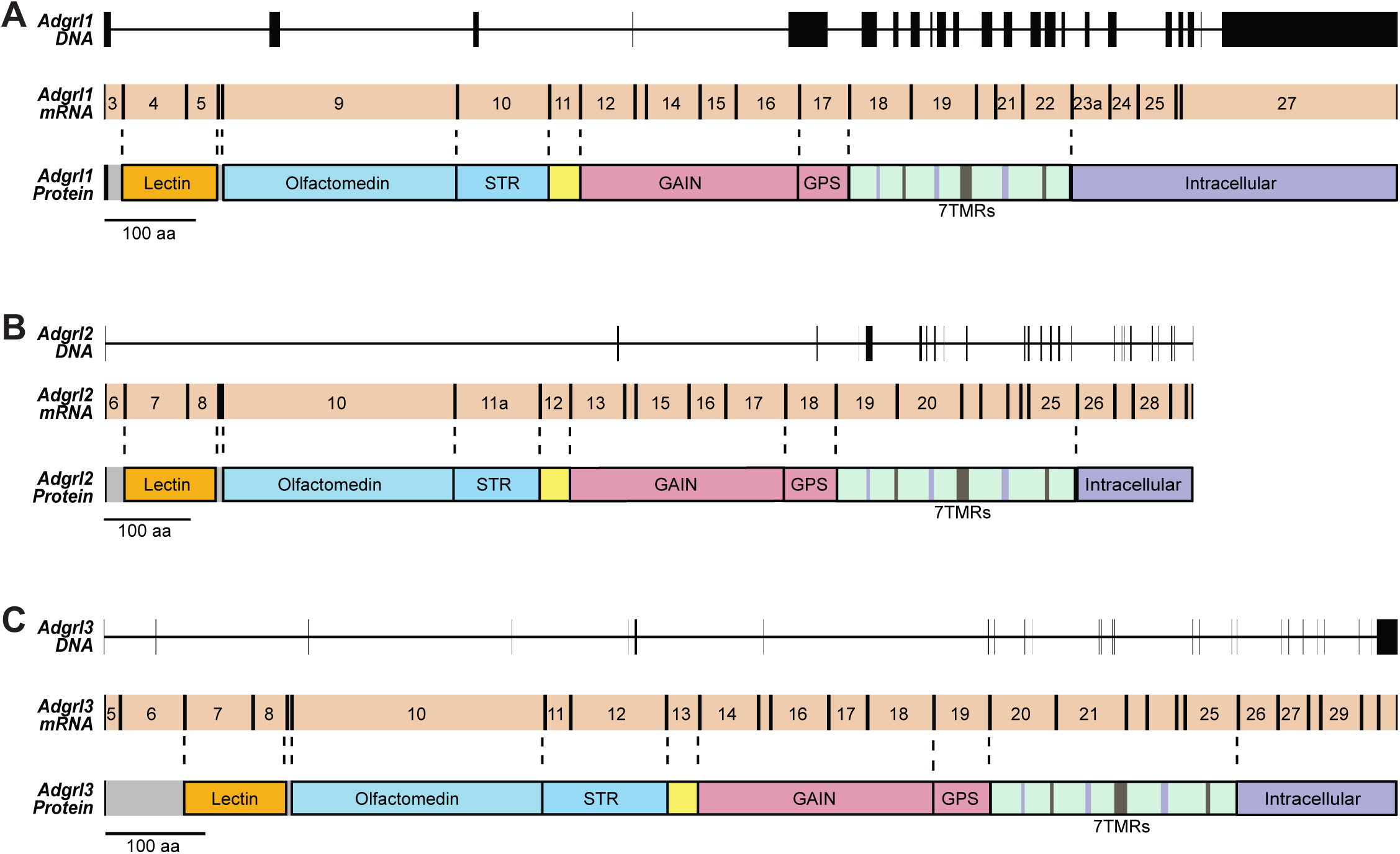
(**A-C**) DNA exon/intron organization, mRNA exon and protein coding alignments for *Adgrl*1 (**A**), *Adgrl2* (**B**), and *Adgrl3* (**C**). For exon number and sequence information, see Supplemental Tables 1-3.

**Supplemental Figure 3.**
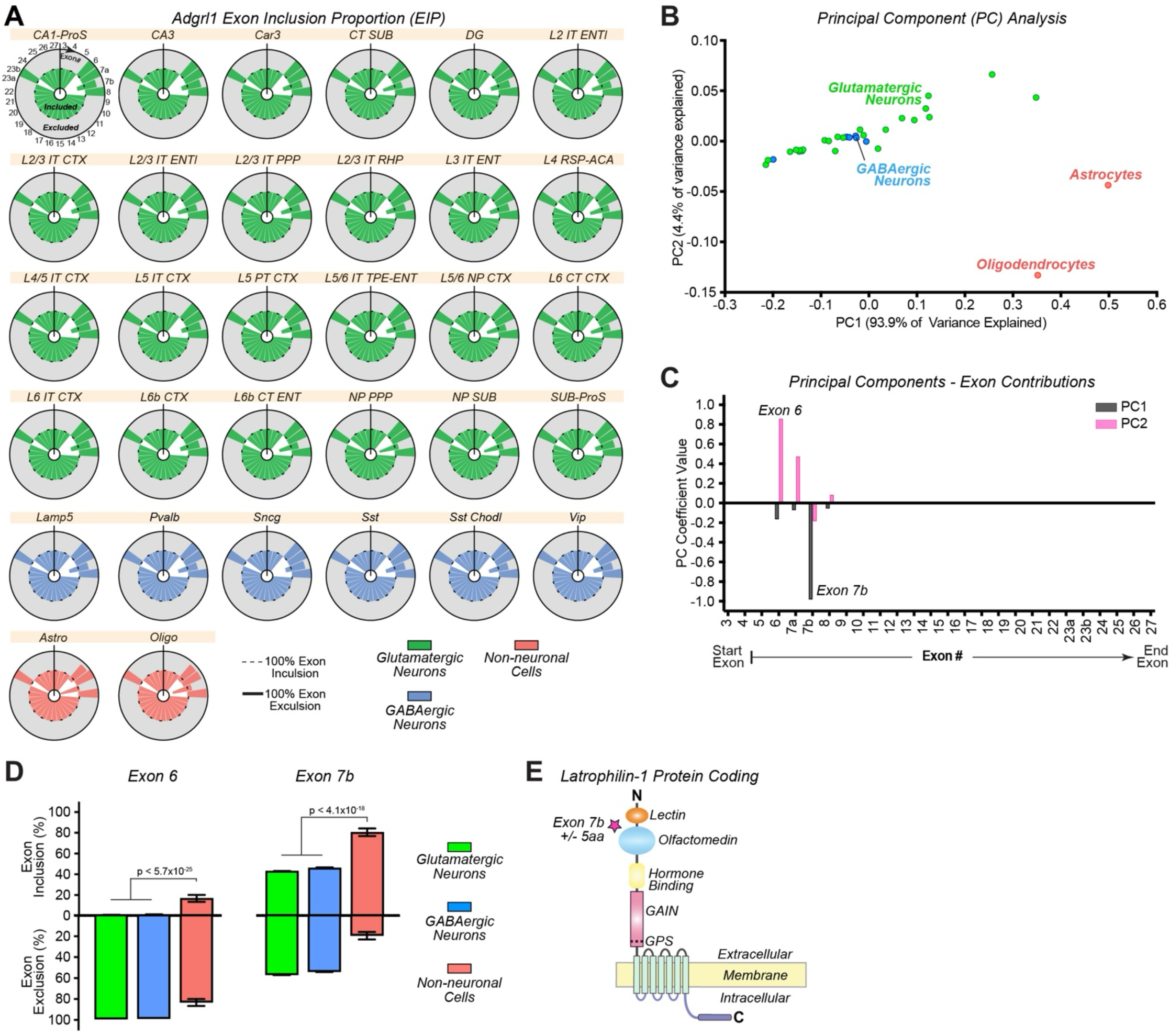
A*d*grl1 exhibits cell type-specific alternative splicing limited to the lectin-olfactomedin domain extracellular region. (**A**) *Adgrl1* splicing profiles for each exon for each cell subclass that expresses *Adgrl1*. Shown are exon inclusion proportion (EIP) circle charts with 5’ start exon indicated, and downstream exons progressing clockwise. Indicated exon is maximally included if graphic slice is confined to interior circle (dashed line), or maximally excluded if slice is restricted to shaded gray region of outer circle (solid line). (**B**) Principal component analysis (PCA) for different cell subtypes based on their alternative splicing patterns. Each point corresponds to a cell subtypes color coded by major classifications including glutamatergic (green), GABAergic (blue), and non-neuronal cells (red). Plot axes represent the first two main components that capture most variance in splicing profiles. (**C**) Summary graph indicating the contribution of individual *Adgrl1* exons to the first two principal components identified in the PCA. (**D**) Summary graphs of exon inclusion proportions (top) and exon exclusion proportions (bottom) for the major alternatively spliced exons (6 and 7b) averaged across all Glutamatergic neurons, GABAergic neurons, and non-neuronal cell classes. Statistical analysis was performed by Dunn’s Test of multiple comparisons, with significance indicated. Only those cells containing transcripts that detected inclusion or exclusion of an exon were included for statistical analysis (n = 35843-35878 Glutamatergic neurons; 13432-13449 GABAergic neurons; 118-122 non-neuronal cells). (**E**) Visualization of *Adgrl1* protein structure. Star indicates the domain region where alternative splice sites would impact protein coding.

**Supplemental Figure 4.**
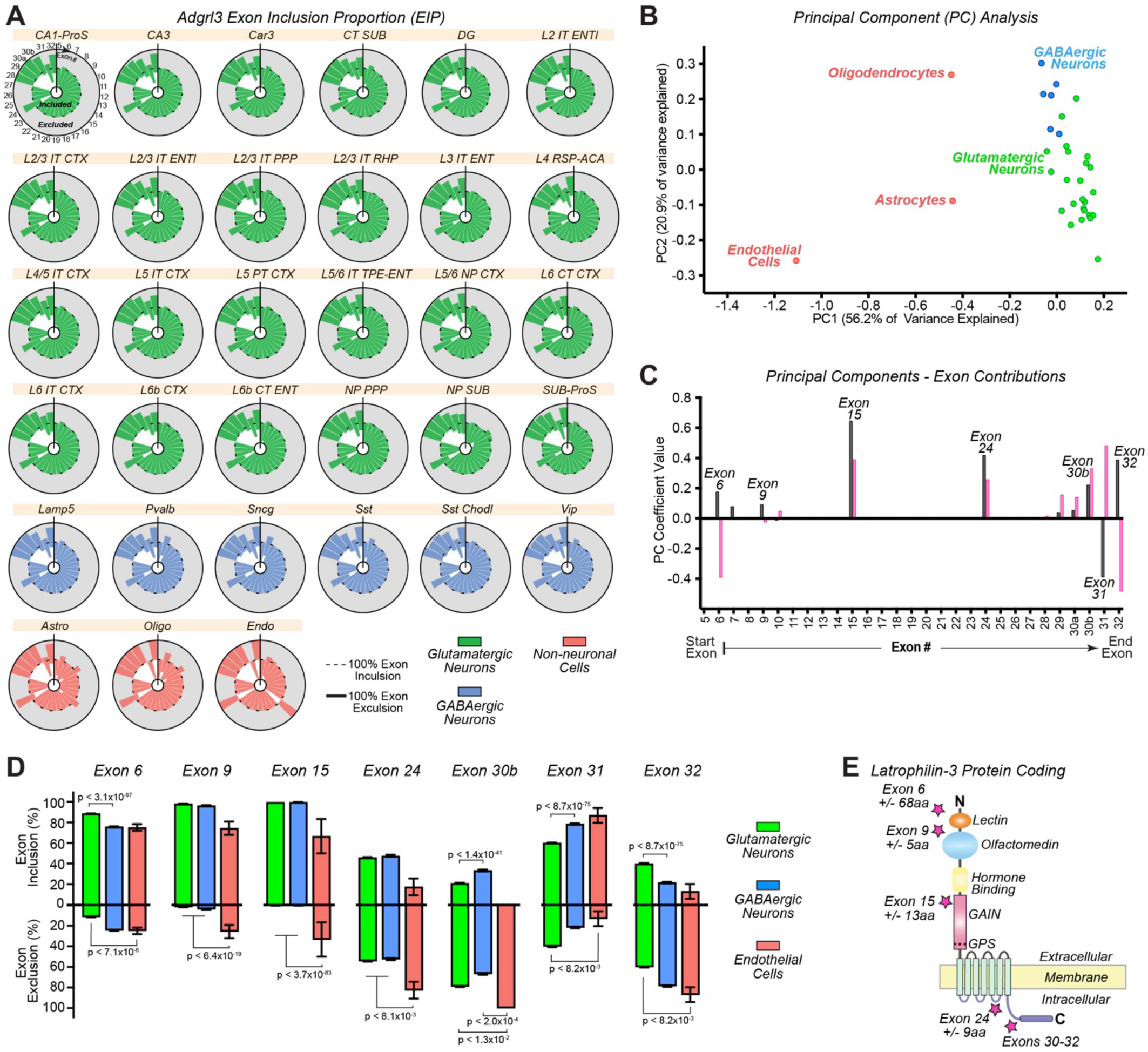
*Adgrl3* is alternatively spliced at five major structural regions. **(A)** *Adgrl3* splicing profiles for each exon for each cell subclass that expresses *Adgrl3*. Shown are exon inclusion proportion (EIP) circle charts with 5’ start exon indicated, and downstream exons progressing clockwise. Indicated exon is maximally included if graphic slice is confined to interior circle (dashed line), or maximally excluded if slice is restricted to shaded gray region of outer circle (solid line). (**B**) Principal component analysis (PCA) for different cell subtypes based on their alternative splicing patterns. Each point corresponds to a cell subtypes color coded by major classifications including glutamatergic (green), GABAergic (blue), and non-neuronal cells (red). Plot axes representing the first two principal components that capture the majority of variance in splicing profiles. (**C**) Summary graph indicating the contribution of individual *Adgrl3* exons to the first two principal components identified in the PCA. (**D**) Summary graphs of exon inclusion proportions (top) and exon exclusion proportions (bottom) for the major alternatively spliced exons (6, 9, 15, 24, and 30-32) averaged across all Glutamatergic neurons, GABAergic neurons, and non-neuronal cell classes. Statistical analysis was performed by Dunn’s Test of multiple comparisons, with significance indicated. Only those cells containing transcripts that detected inclusion or exclusion of an exon were included for statistical analysis (n = 4115-14237 Glutamatergic neurons; 1581-4682 GABAergic neurons; 9-212 non-neuronal cells). (**E**) Visualization of *Adgrl2* protein structure. Stars indicate the domain regions where exons 6, 9, 15, 24, and 30-32 alternative splice sites would impact protein coding.

**Supplemental Figure 5.**
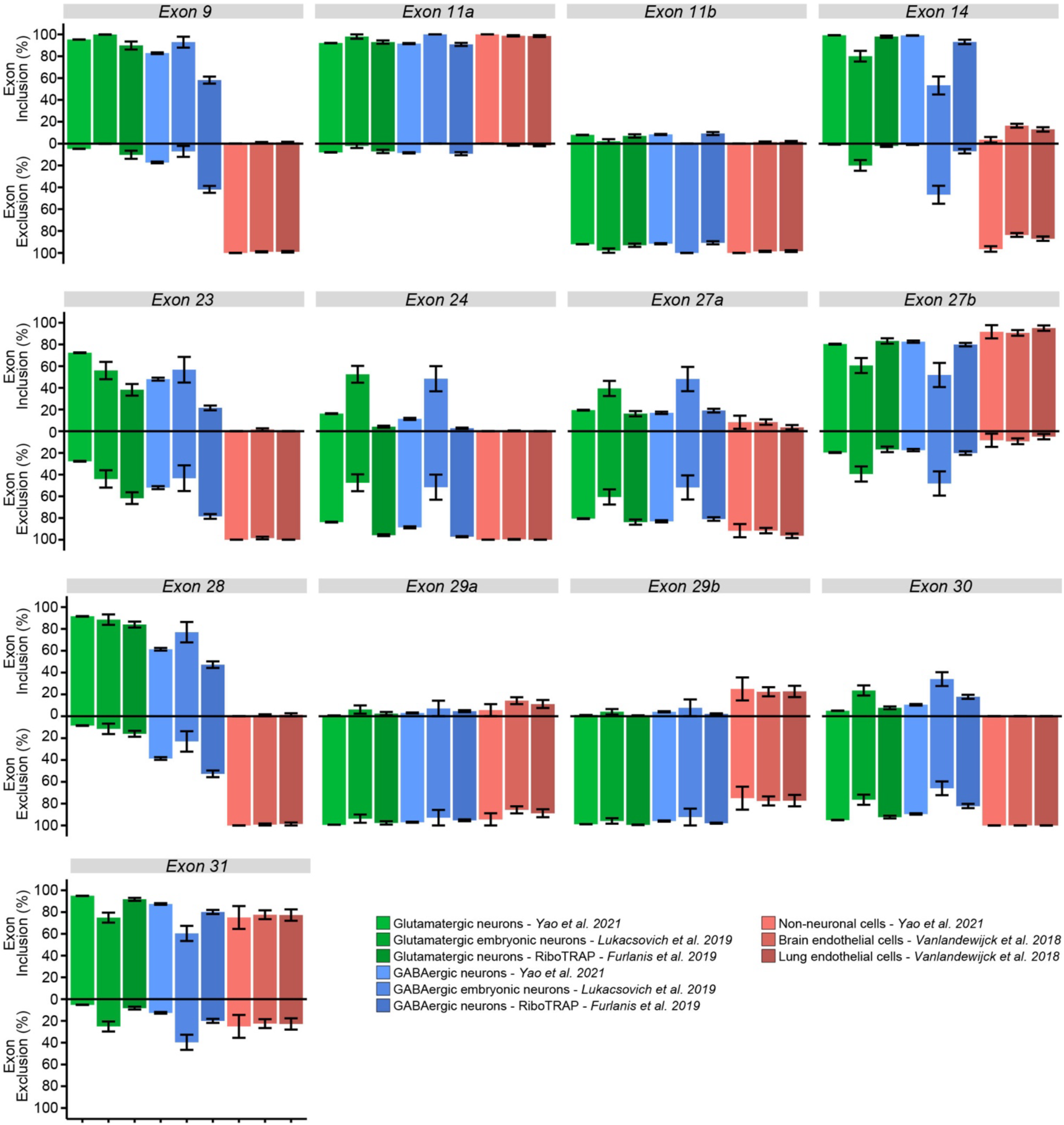
*Adgrl2* alternative splicing. Summary graphs of exon inclusion proportions (EIP) and for alternatively spliced exons of *Adgrl2* (9, 11a, 11b, 14, 23, 24, 27a, 27b, 28, 29a, 29b, 30, 31) averaged across all Glutamatergic neurons, GABAergic neurons, non-neuronal cells, and endothelial specific RNAseq datasets from four independent studies (Furlanis et al., 2019; Lukacsovich et al., 2019; Vanlandewijck et al., 2018; Yao et al., 2021).

**SUPPLEMENTAL TABLE 1.**
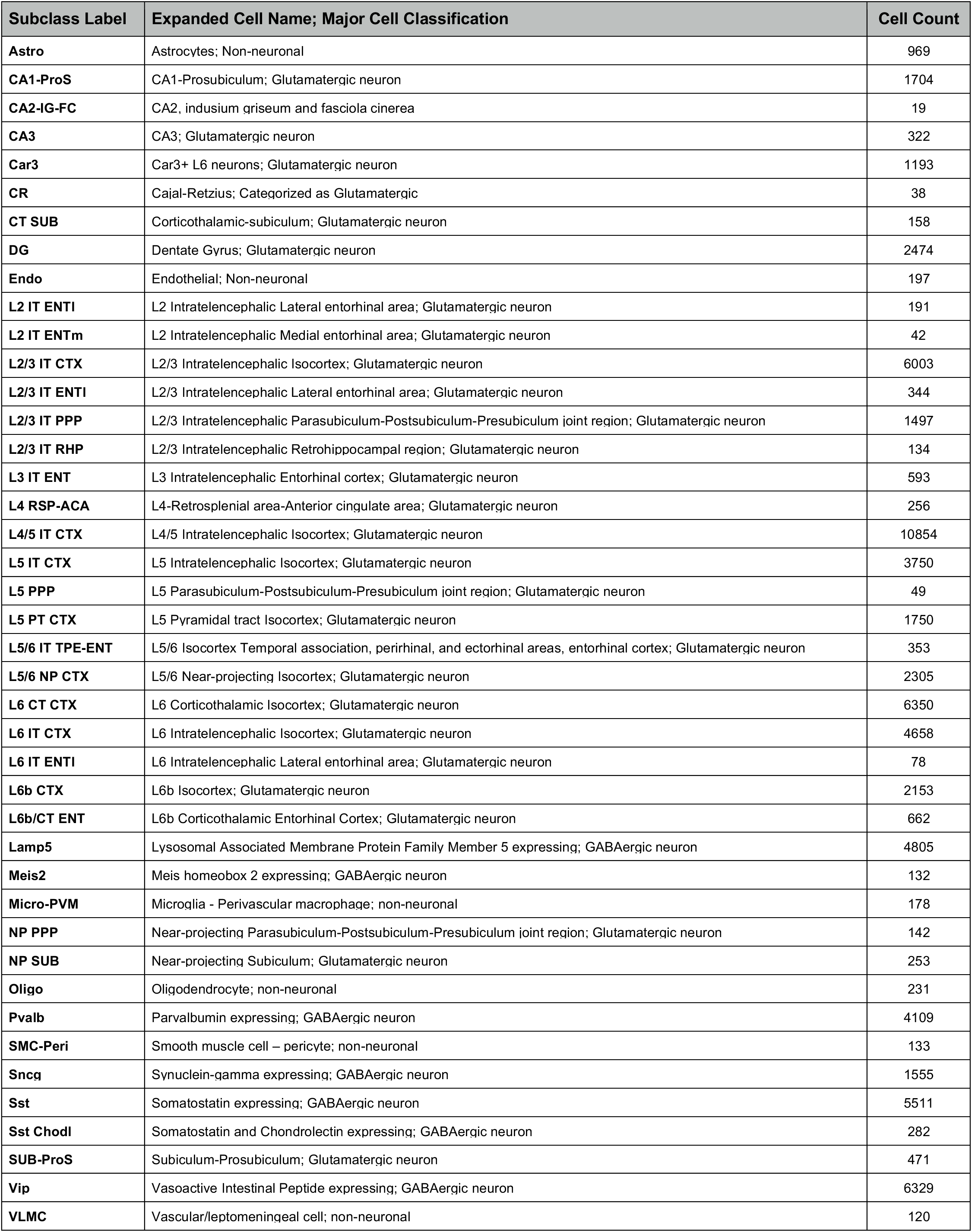

**SUPPLEMENTAL TABLE 2.**
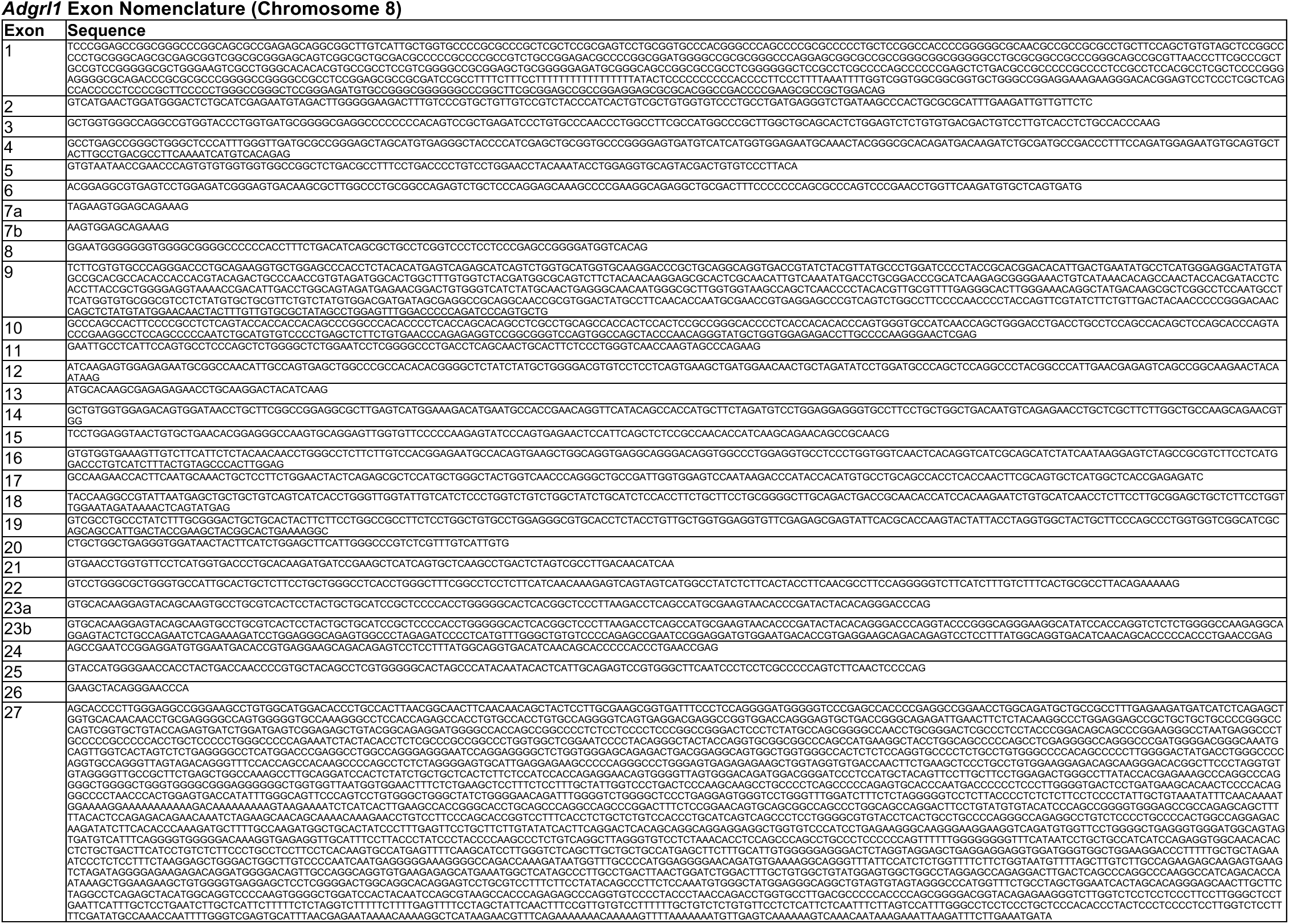

**SUPPLEMENTAL TABLE 3.**
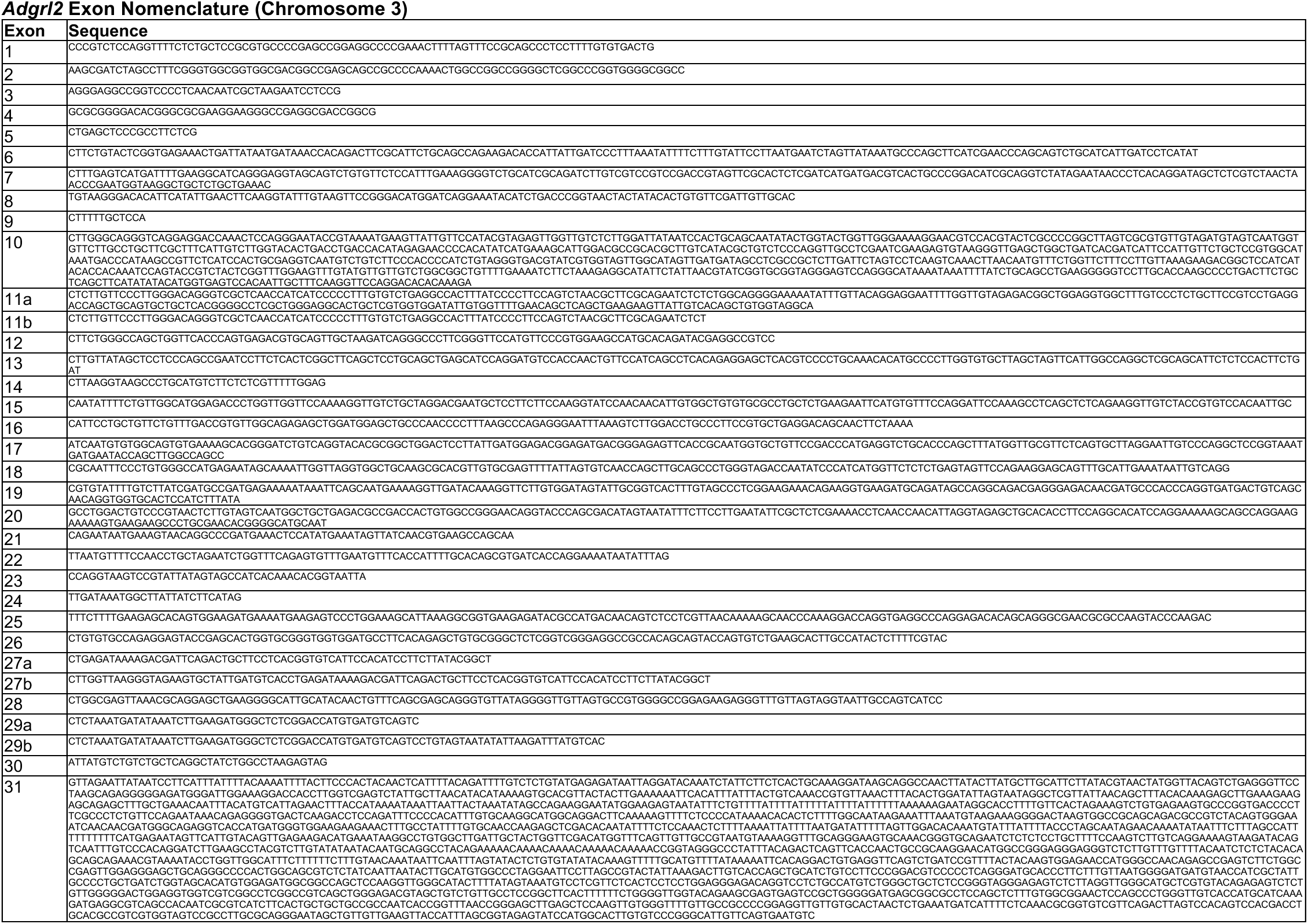

**SUPPLEMENTAL TABLE 4.**
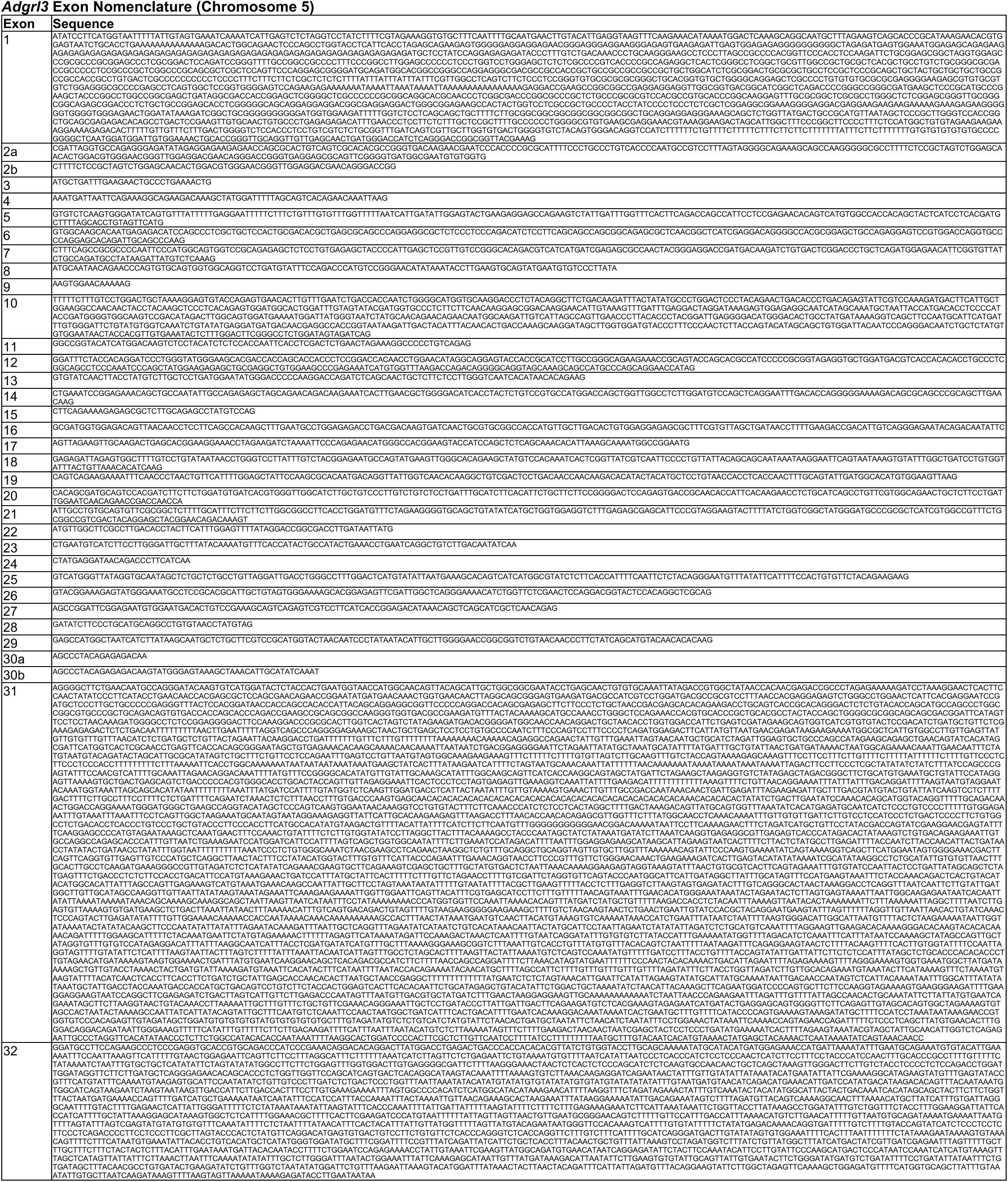

## Notes

### Competing Interest Statement

The authors have declared no competing interest.

